# Suction feeding of West African lungfish (*Protopterus annectens*): An XROMM analysis of jaw mechanics, cranial kinesis, and hyoid mobility

**DOI:** 10.1101/2022.05.30.493759

**Authors:** Samantha M. Gartner, Katrina R. Whitlow, J.D. Laurence-Chasen, Elska B. Kaczmarek, Michael C. Granatosky, Callum F. Ross, Mark W. Westneat

**Affiliations:** Department of Organismal Biology and Anatomy, University of Chicago, 1027 East 57^th^ St, Chicago, IL 60637; Department of Ecology, Evolution, and Organismal Biology, Brown University, 80 Waterman St., Providence RI 02912; Department of Anatomy, New York Institute of Technology College of Osteopathic Medicine, 100 Northern Blvd, Old Westbury, NY 11568

**Author notes:** Corresponding Author Information: Samantha M. Gartner, 1025 E 57th St, Culver 105, Westneat Lab.

**Keywords:** sarcopterygian, biomechanics, x-ray reconstruction of moving morphology, feeding kinematics

## Abstract

Suction feeding in fishes is characterized by rapid cranial movements, but extant lungfishes (Sarcopterygii: Dipnoi) exhibit a reduced number and mobility of cranial bones relative to actinopterygian fishes. Despite fusion of cranial elements, lungfishes are proficient at suction feeding, though the impacts of novel cranial morphology and reduced cranial kinesis on feeding remain poorly understood. We used X-ray Reconstruction of Moving Morphology (XROMM) to study the kinematics of seven mobile skeletal elements (neurocranium, upper jaw, lower jaw, tongue, ceratohyal, clavicle, and cranial rib) and two muscles (costoclavicular portion of the hypaxialis and rectus cervicis) during the feeding strikes of West African lungfish (*Protopterus annectens*). We found that feeding by *P. annectens* on non-evasive prey is relatively slow, with a mean time to peak gape of 273 ms. Lower jaw depression and clavicular rotation were hinge-like, with one degree of freedom, but the ceratohyals rotated in a complex motion involving depression and long-axis rotation. We quantified the relative contributions to oral cavity volume change (RCVC) and found that oral cavity expansion is created primarily by ceratohyal and clavicle motion. *P. annectens* suction feeds relatively slowly but successfully through muscle shortening of hypaxial and rectus cervicis muscles contributing to hyoid mobility.

**Summary Statement:** Three-dimensional hyoid movements and clavicle retraction generate suction during the relatively slow, but successful, feeding strikes of the West African lungfish (*Protopterus annectens*).

## Introduction

Suction feeding is the predominant feeding strategy in fishes and is one of the most well-studied behaviors performed by a highly complex and mobile musculoskeletal system. The majority of the literature on suction feeding in fishes has focused on cranial kinesis in actinopterygians (e.g. Day et al., 2015; Lauder, 1982; Liem, 1980; Wainwright et al., 2015; Westneat, 2005) or chondrichthyans (Motta and Wilga, 2001; Wilga et al., 2007, 2000). The kinetic skulls of suction feeding actinopterygians enable considerable expansion of the oral cavity, allowing some fish species to eat very large prey (Pietsch and Grobecker, 1990) or to extend their jaws anteriorly to capture elusive prey (Westneat and Wainwright, 1989). Additionally, the relative timing of skeletal movements during suction feeding is similar across species: cranial elevation and jaw depression precede hyoid depression, opercular flaring, and pectoral girdle retraction, a pattern that is pivotal for creating suction (Holzman et al., 2007; Whitlow et al., 2022). These skull movements occur in an anterior-to-posterior wave of motion that pulls a volume of water, and the prey, into the buccal cavity during the suction feeding strike (Bishop et al., 2008; Ferry et al., 2015). Suction feeding also typically occurs rapidly, with jaw opening and closing occurring within a relatively short time period, ranging from 1-5 ms in seahorses and pipefishes (Bergert and Wainwright, 1997; Van Wassenbergh et al., 2009) up to a more typical duration of 50-100 ms in fishes such as catfishes (Olsen et al., 2019), wrasses (Westneat, 1990, 1994) and basses (Camp and Brainerd, 2014; Wainwright et al., 2007).

The feeding mechanics of living sarcopterygian fishes such as coelacanths and lungfishes is less well understood than in actinopterygians (Bemis, 1986; Bemis and Lauder, 1986; Dutel et al., 2015b, 2015a). Lungfish feeding kinematics are known primarily from the work of Bemis and Lauder (1986) on *Lepidosiren paradoxa*, the South American lungfish, highlighting the need for studies of cranial function in the other lungfish genera *Neoceratodus* and *Protopterus*. With a strategy similar to ray-finned fishes, the South American lungfish employs an anterior-to-posterior wave of motion during suction feeding (Bemis and Lauder, 1986). However, suction feeding is performed without jaw protrusion, and the duration of the suction strike in *L. paradoxa* is relatively high, often lasting over 300 milliseconds (Bemis and Lauder, 1986). Also, the role of internal structures such as the ceratohyal, clavicle and tongue in lungfishes in buccal expansion is largely unknown, although Bemis and Lauder (1986) measured the ventral extent of ceratohyal depression and suggested that the fleshy tongue “pad” was passive, moving in concert with ceratohyal motion.

Suction feeding in lungfishes also differs from suction feeding in actinopterygian fishes in that lungfishes use a modified set of skull elements and generate suction over a longer period of time (Bemis and Lauder, 1986). Lungfishes possess more bony elements than early tetrapods, but fewer than actinopterygian fishes (Fig. 1; Bemis and Lauder, 1986; Clack et al., 2016; Criswell, 2015; Heiss et al., 2013). The pterygoid bone and prearticular bone are each fused into two large tooth plates that compose the upper and lower jaw, respectively (Fig. 1, Fig. 2A). The number of skeletal elements in their opercular series and hyoid apparatus is also reduced, with the ceratohyal being the only skeletal element of the hyoid bar (Fig. 1).

**Fig. 1.**
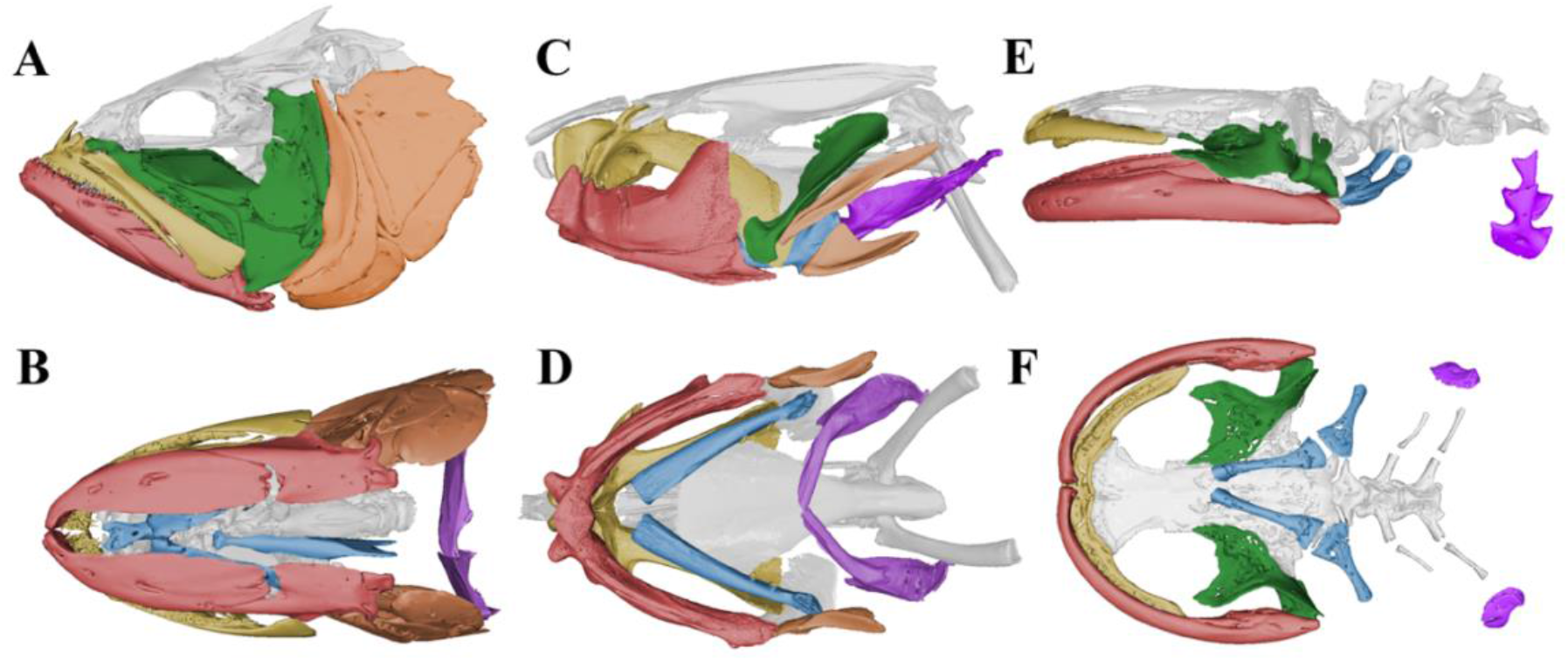
Comparative skull morphology of suction feeding vertebrates. (A) Lateral view of the skull anatomy of a largemouth bass, *Micropterus salmoides*, and (B) same *M. salmoides* skull in ventral view. (C) Skull of the West African lungfish, *Protopterus annectens* in lateral view and (D) in ventral view showing the large hyoid apparatus. (E) Skull anatomy of a Japanese giant salamander, *Andrias japonicus* also in (F) ventral view. Coloration represents similar skeletal structures across the three species; red—lower jaw, orange—opercular series, yellow—upper jaw, green—suspensorium, blue—hyoid apparatus, purple—pectoral girdle, gray—neurocranial elements. Largemouth bass mesh from Morphosource (ark:/87602/m4/M26211; uf:fish:34881.1). Japanese giant salamander mesh from Morphosource (ark:/87602/m4/M162014; fmnh:amphibians and reptiles:31536).

**Fig. 2.**
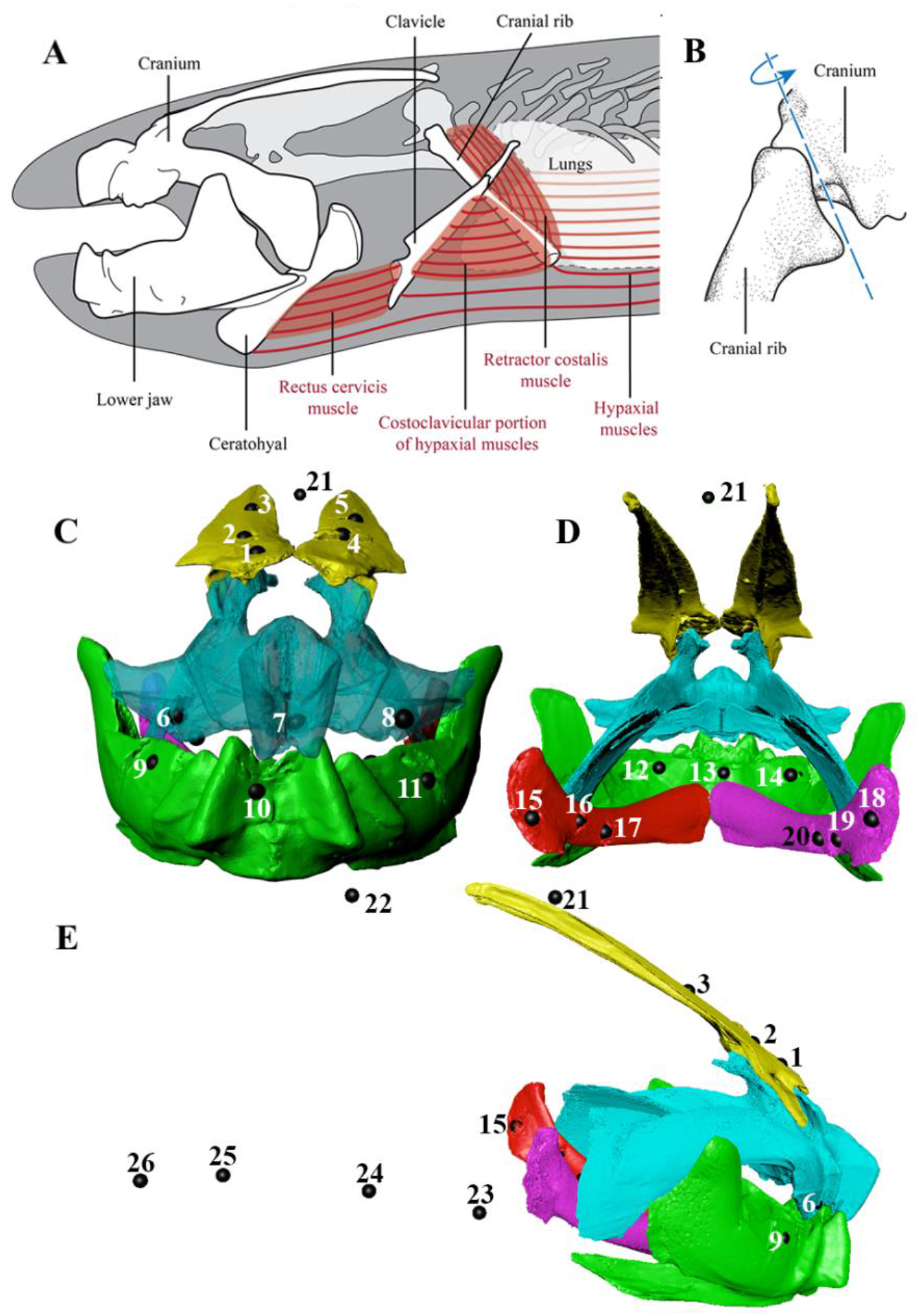
Cranial musculoskeletal anatomy of West African lungfish, *Protopterus annectens* and general bead placement in the lungfish skull and body plane in lateral view. (A) Lateral view of the animated bones (labeled), rectus cervicis muscle, costoclavicular portion of the hypaxial muscles (CCH), retractor costalis muscle, and hypaxial muscles. Note that, for clarity, not all bones of the head are shown. (B) Posterolateral view of the costo-cranial joint, which connects the cranial rib and the exoccipital bone of the cranium. Primary axis of motion (based on morphology) is shown in blue. Modified from Kaczmarek et al. (2022). (C) displays the frontal view, (D) caudal view, and (E) lateral view of the total beads placed in each individual. Three beads were placed in each of the following bones: upper jaw, lower jaw, and right and left ceratohyal. Five beads were placed in the neurocranium. One to two beads were placed in the dorsal body plane and four beads were placed in the ventral body plane. Points 12-14 in panel E represent the tongue markers.

Lungfishes are the closest living relative to tetrapods (Amemiya et al., 2013), making them useful for understanding the traits and behaviors fishes may have employed prior to the water-to-land transition. The overall trend in lungfish skull evolution has been to reduce their skeletal elements and evolve robust jaws to crush their food (Fig. 1, Fig. 2A; Clack et al., 2016; Clack and Ahlberg, 2016), suggesting a trade-off between high cranial mobility for suction performance and increased bite force during processing. Despite the loss and fusion of bones in the skull and the long duration of the suction strikes, suction feeding in the South American lungfish (*Lepidosiren paradoxa*) is frequently successful (Bemis and Lauder, 1986). Given that the magnitude of sub-ambient pressure and resultant velocity of water depend on the rate of buccal expansion, how do lungfishes generate the suction flow required to successfully capture prey? We hypothesized that the long strike duration in lungfishes is combined with large ceratohyal depression and clavicular retraction, leading to buccal cavity expansion, and that these motions enable successful suction strikes by mobilizing a volume of water over a longer temporal duration.

To test this hypothesis, we investigated the feeding biomechanics of the West African lungfish, *Protopterus annectens*, using biplanar videoradiography and the X-ray Reconstruction of Moving Morphology workflow (XROMM; Brainerd et al., 2010). We quantified the three-dimensional rotations and translations of key elements in the feeding system of *P. annectens*, including internal elements such as the ceratohyal, clavicle and tongue, which are not visible to standard light cameras. We used a recently developed measurement (Whitlow et al. 2022) for quantifying the relative contribution of each cranial bone to the volume change of the oral cavity (RCVC) to assess the roles of individual skull components and test the prediction that complex hyoid motions are the major contributor to suction volume change. We measured the shortening of the rectus cervicis and the costoclavicular portion of the hypaxial muscles, two muscles that are thought to play a role in buccal cavity expansion via the transfer of force and motion to the clavicle and hyoid. Lastly, we quantified the movement of the fleshy tongue pad to evaluate its role in lungfish suction feeding. By exploring the detailed suction feeding kinematics of *Protopterus* and comparing it to that of *Lepidosiren*, actinopterygians, and suction feeding tetrapods, we develop a comparative framework for examining lungfish feeding mechanics across a diversity of suction feeding vertebrates.

## Results

The results of this study show that cranial, hyoid and pectoral kinematics enable lungfishes to perform one of the slowest suction feeding strikes among aquatic organisms. We found that to capture non-evasive prey, *Protopterus annectens* used a suction feeding strike that took 273.1 ± 3.01 ms (mean ± s.e.m.) to reach peak gape, with a strike duration of about 465 ms (Table 1). Cranial elevation was relatively small, with a maximum rotation of about 4.4 deg, whereas the lower jaw depressed up to −39 deg from rest (Fig. 4, Fig. 5). The prey moved up to an average speed of 63.3 ± 3.76 cm s^-1^ into the mouth and reached its maximum velocity around the time of peak gape (Fig. 6). Buccal volume change was small, increasing by an average of 3.8 ± 0.074 cm^3^ (Fig. 7). Cranial kinematics demonstrated an anterior-to-posterior wave of movement, involving sequential rotations of the neurocranium, lower jaw, ceratohyal, clavicle, and cranial rib, as well as substantial long-axis rotation of the ceratohyal. (Fig. 5, Fig. 6, Fig. 8B, Table S3). The rectus cervicis and costoclavicular portions of the hypaxial muscles consistently shortened during strikes, contributing to buccal expansion (Fig. 8A).

**Fig. 3.**
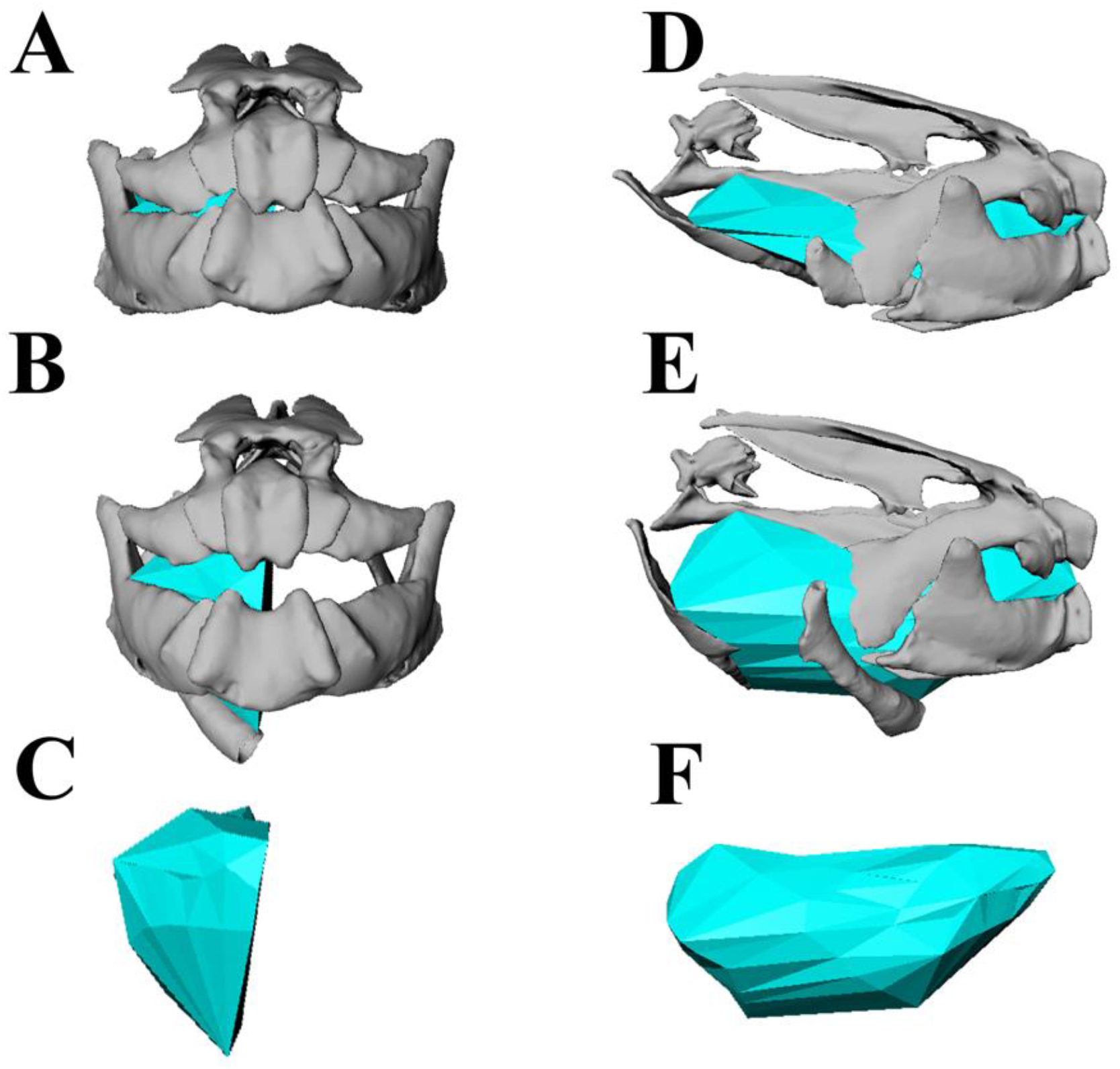
Dynamic digital endocast for the oral cavity volume. Skeletal animations during volume expansion for a suction feeding event. Rostral and lateral view of an animation at the start of the strike (A,D) and at the peak oral cavity volume (B,E). Digital endocast animations shown in a rostral and lateral view are at peak oral cavity volume (C, F).

**Fig. 4.**
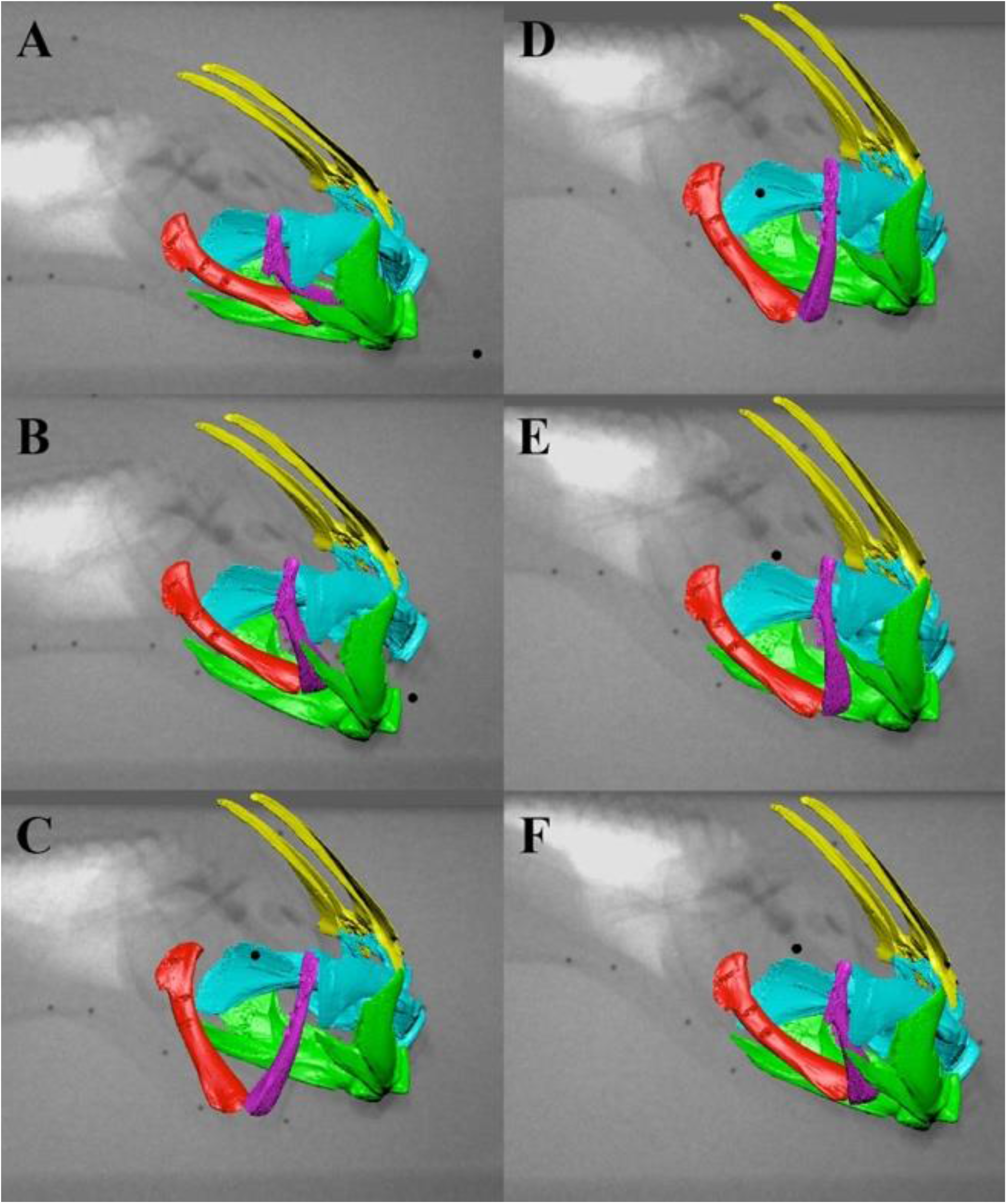
Representative suction feeding sequence animation with bone models and X-ray images. The first three panels are during buccal expansion. A is the resting phase just before the strike begins. B is the time at peak lower jaw depression and C is the time at peak ceratohyal depression. D, E, F represent time points during the buccal compression phase.

**Fig. 5.**
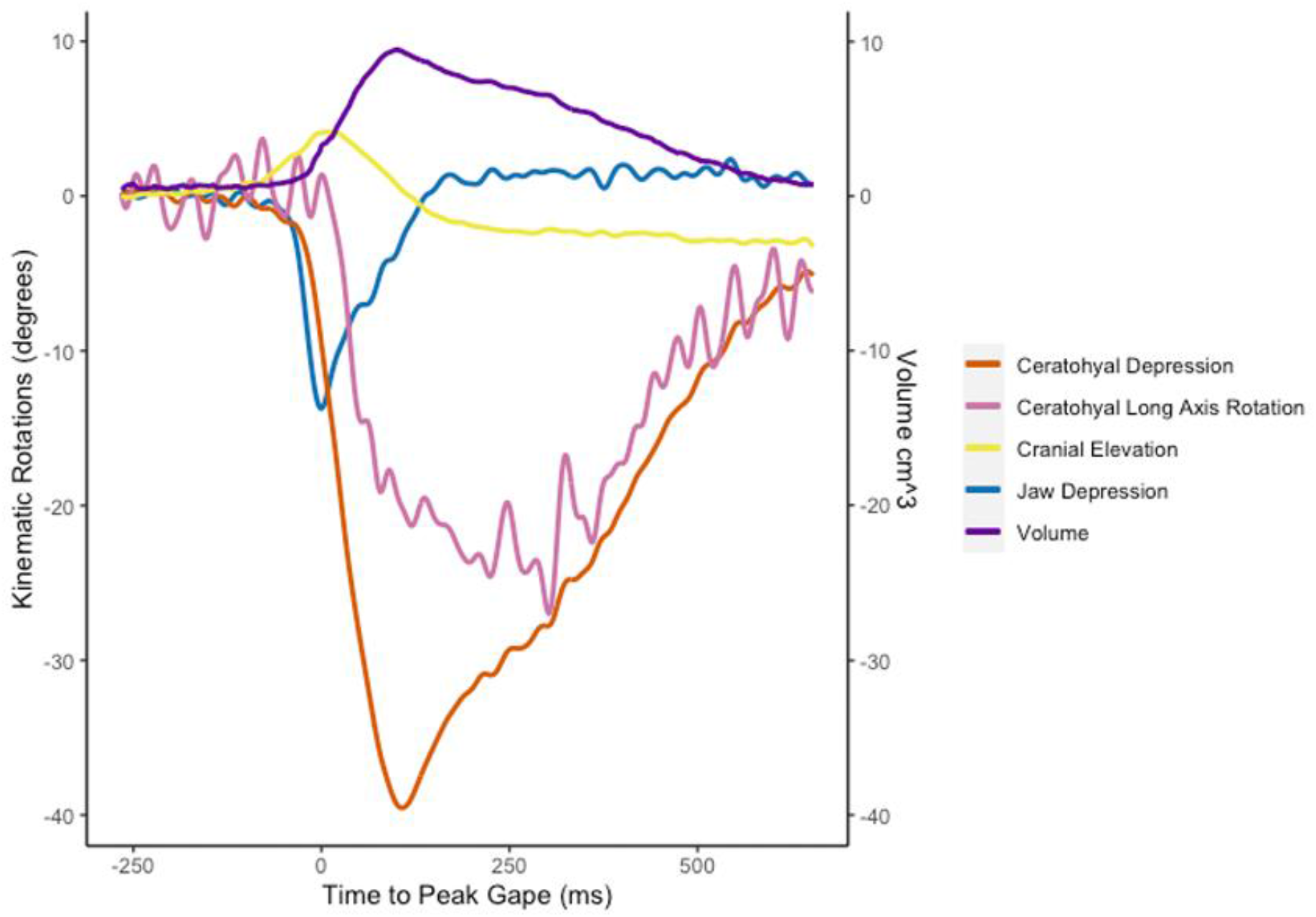
Skeletal kinematics and oral cavity volume from a representative feeding trial. Negative rotation indicates depression of the cranium, lower jaw, and ceratohyal and internal long-axis rotation of the ceratohyal. Ceratohyal depression and long axis rotation and jaw depression were measured relative to the neurocranium while cranial elevation was measured relative to the body plane. See Methods and Materials for more in depth descriptions of each measurement.

**Fig. 6.**
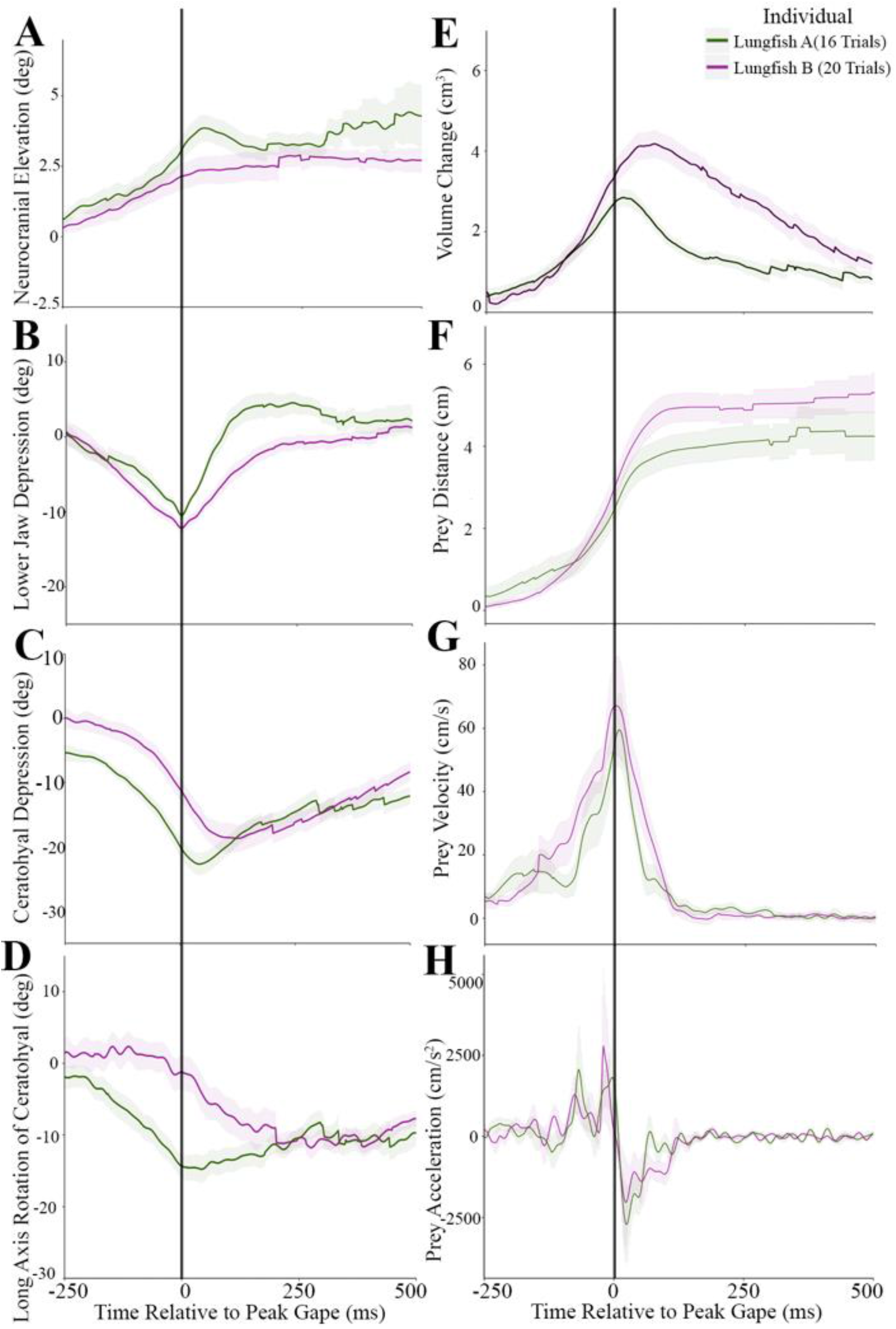
Trial-average plots for skeletal kinematics, oral volume change, and prey motion. (A) Cranial elevation, (B) lower jaw depression, (C) ceratohyal depression, (D) long axis rotation of the ceratohyal, (E) volume change, (F) prey distance, (G) prey velocity, (H) prey acceleration. The vertical black line indicates time = 0 or the time at peak gape. Shaded regions show the standard error around the mean.

**Fig. 7.**
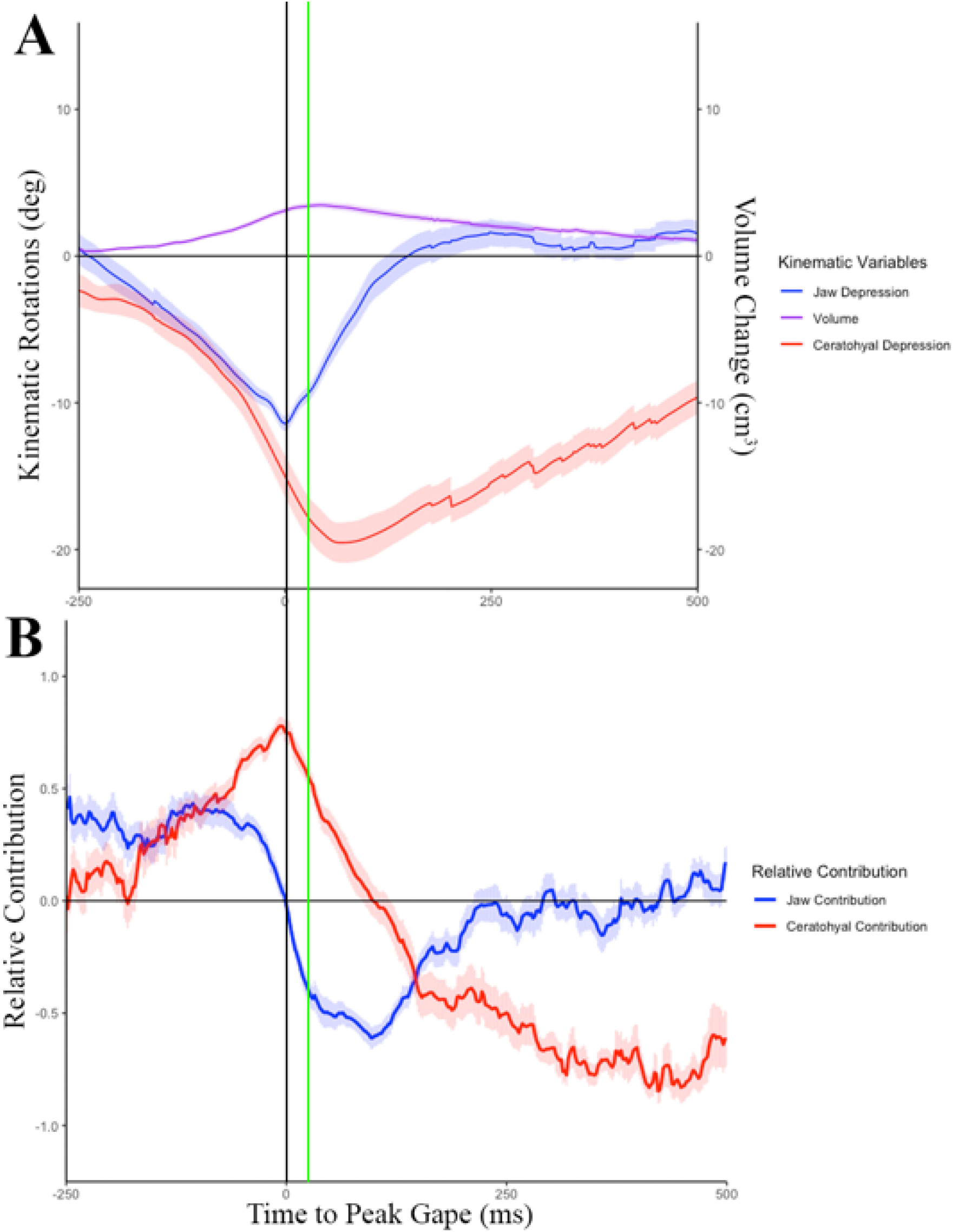
Skeletal kinematics and the relative contribution to volume change. Graphs of average lower jaw depression, volume change, and ceratohyal depression (A) and the relative contribution of the lower jaw and ceratohyal to oral cavity volume change (B) averaged across the two individuals (n = 36 trials). The vertical black line indicates time at peak gape. The vertical green line is indicating time at peak volume.

**Fig. 8.**
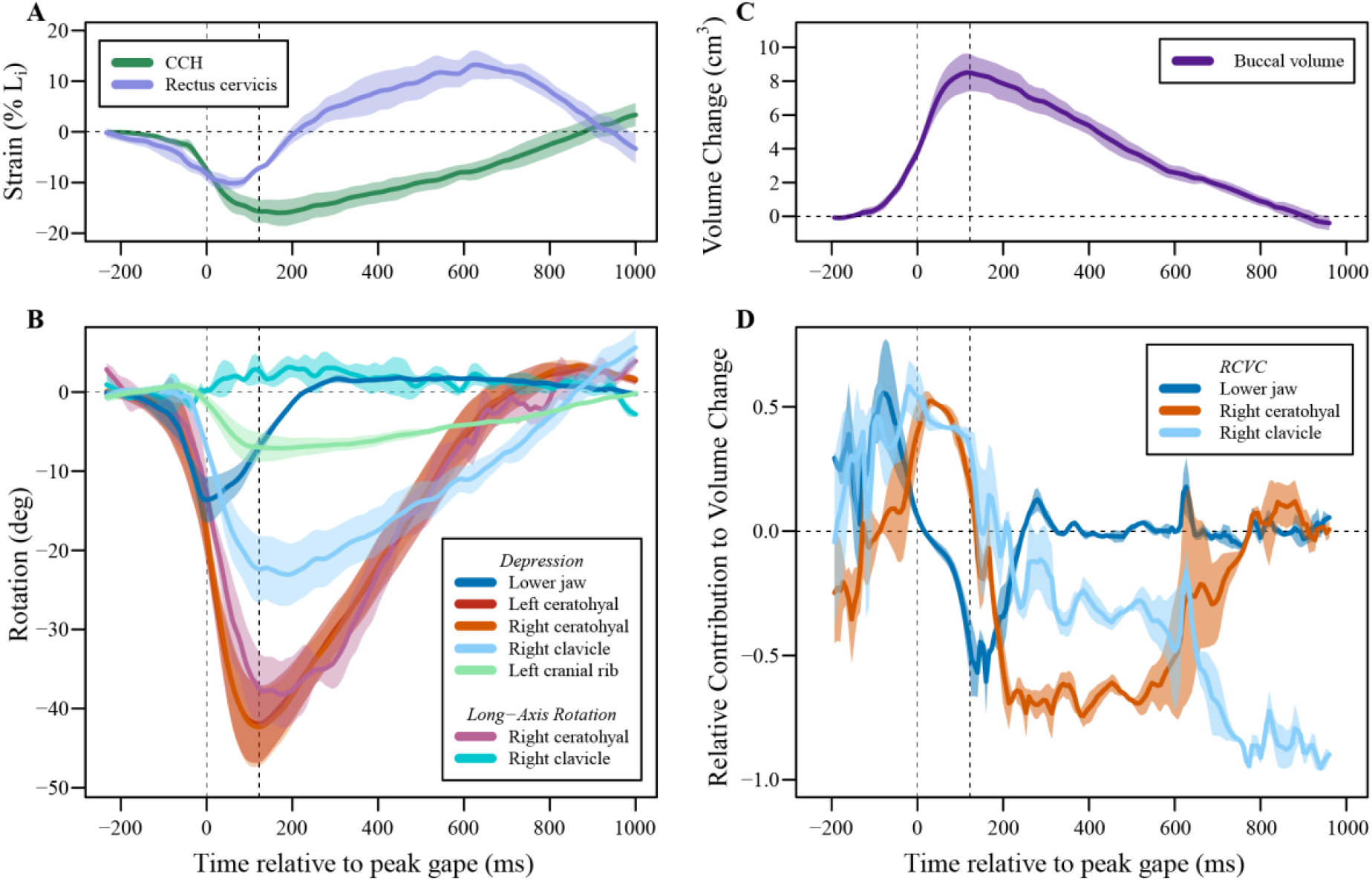
Muscle shortening, skeletal kinematics, buccal volume, and relative contribution to volume change for three strikes from lungfish B in which the clavicles and cranial rib were marked. All values are shown as mean ± s.e.m. Left dashed line indicates the time of peak jaw depression, and right dashed line indicates the time of peak ceratohyal depression. (A) Strain of the rectus cervicis and CCH, calculated as percent change in length relative to the initial length (Li). (B) Z-axis rotation (depression) of the lower jaw, ceratohyals, right clavicle, and left cranial rib, and x-axis (long-axis) rotation of the right ceratohyal and right clavicle. Negative values represent depression or internal long-axis rotation, and positive values represent elevation or external long-axis rotation. (C) Change in buccal volume, normalized to initial volume. (D) Relative contributions to volume change of the lower jaw, right ceratohyal, and right clavicle. Positive values indicate that the motion of the bone is contributing to expansion of the buccal volume, and negative values indicate that the motion of the bone is contributing to contraction of the buccal volume.

**Table 1.** Average kinematic variables across lungfish feeding strikes. Timings are relative to the start of the suction feeding strike. Values are given as mean ± SE. Note that strike duration was measured as the time from onset of jaw opening to end of jaw closing.

| Variables | Lungfish A<br>(n = 16) | Lungfish B<br>(n = 20) | Combined Average<br>(n = 36) |
| --- | --- | --- | --- |
| Peak Jaw Depression (deg) | -10.5 ± 0.65 | -12.2 ± 0.72 | -11.3 ± 0.85 |
| Time to Peak Gape (ms) | 276.1 ± 17.76 | 270.1 ± 24.22 | 273.1 ± 3.01 |
| Peak Ceratohyal Depression (deg) | -22.7 ± 1.74 | -18.7 ± 2.11 | -20.7 ± 2.01 |
| Time to Peak Ceratohyal Depression (ms) | 337.1 ± 20.35 | 397.5 ± 21.74 | 367.3 ± 30.19 |
| Peak Ceratohyal Long Axis Rotation | -14.8 ± 1.88 | -11.8 ± 1.52 | -13.3 ± 1.50 |
| Peak Volume (cm<sup>3</sup>) | 2.9 ± 0.05 | 4.5 ± 0.12 | 3.8 ± 0.07 |
| Time to Peak Volume (ms) | 310.0 ± 11.76 | 359.7 ± 14.45 | 334.8 ± 27.86 |
| Cranial Elevation (deg) | 4.4 ± 1.09 | 2.9 ± 0.39 | 3.8 ± 0.76 |
| Time to Peak Cranial Elevation (ms) | 46.0 ± 0.47 | 48.0 ± 0.49 | 47.0 ± 0.47 |
| Maximum Prey Distance (cm) | 4.4 ± 0.50 | 5.3 ± 0.42 | 4.9 ± 0.45 |
| Time to Maximum Prey Distance | 564.1 ± 42.67 | 596.9 ± 55.52 | 580.5 ± 19.4 |
| Maximum Prey Velocity (cm/s) | 59.6 ± 11.7 | 67.1 ± 15.58 | 63.3 ± 3.76 |
| Maximum Prey Acceleration (cm/s<sup>2</sup>) | 2066.7 ± 1349.49 | 2798.7 ± 2351.2 | 2432.7 ± 366.03 |
| Strike Duration (ms) | 447.1 ± 125.26 | 484.0 ± 164.13 | 465.6 ± 26.07 |

**Table 1. Average kinematic variables across lungfish feeding strikes. Timings are relative to the start of the suction feeding strike. Values are given as mean ± SE. Note that strike duration was measured as the time from onset of jaw opening to end of jaw closing.**
| **Variables** | **Lungfish A**<br>**(n = 16)** | **Lungfish B**<br>**(n = 20)** | **Combined**<br>**Average**<br>**(n = 36)** |
| --- | --- | --- | --- |
| Peak Jaw Depression (deg) | -10.5 ± 0.65 | -12.2 ± 0.72 | -11.3 ± 0.85 |
| Time to Peak Gape (ms) | 276.1 ± 17.76 | 270.1 ± 24.22 | 273.1 ± 3.01 |
| Peak Ceratohyal Depression (deg) | -22.7 ± 1.74 | -18.7 ± 2.11 | -20.7 ± 2.01 |
| Time to Peak Ceratohyal Depression (ms) | 337.1 ± 20.35 | 397.5 ± 21.74 | 367.3 ± 30.19 |
| Peak Ceratohyal Long Axis Rotation | -14.8 ± 1.88 | -11.8 ± 1.52 | -13.3 ± 1.50 |
| Peak Volume (cm<sup>3</sup>) | 2.9 ± 0.05 | 4.5 ± 0.12 | 3.8 ± 0.07 |
| Time to Peak Volume (ms) | 310.0 ± 11.76 | 359.7 ± 14.45 | 334.8 ± 27.86 |
| Cranial Elevation (deg) | 4.4 ± 1.09 | 2.9 ± 0.39 | 3.8 ± 0.76 |
| Time to Peak Cranial Elevation (ms) | 46.0 ± 0.47 | 48.0 ± 0.49 | 47.0 ± 0.47 |
| Maximum Prey Distance (cm) | 4.4 ± 0.50 | 5.3 ± 0.42 | 4.9 ± 0.45 |
| Time to Maximum Prey Distance | 564.1 ± 42.67 | 596.9 ± 55.52 | 580.5 ± 19.4 |
| Maximum Prey Velocity (cm/s) | 59.6 ± 11.7 | 67.1 ± 15.58 | 63.3 ± 3.76 |
| Maximum Prey Acceleration (cm/s<sup>2</sup>) | 2066.7 ± 1349.49 | 2798.7 ± 2351.2 | 2432.7 ± 366.03 |
| Strike Duration (ms) | 447.1 ± 125.26 | 484.0 ± 164.13 | 465.6 ± 26.07 |

### Suction feeding kinematics

The rotations of the neurocranium, lower jaw, and right ceratohyal reveal the kinematic pattern of cranial elements in *Protopterus* strikes (Fig. 4, Fig. 5). Neurocranial elevation was small and highly variable within individuals. The neurocranium rotated positively (elevated) about the transverse axis (z-axis) an average of 3.8 ± 0.76 deg (Fig. 6A), reaching its peak elevation 47.0 ± 0.47 ms after peak gape (Table 1). Although neurocranial elevation occurred in the majority of trials, neurocranial depression was occasionally observed immediately before neurocranial elevation (Fig. S4). Peak neurocranial depression ranged from about −2 to −4 deg.

Strike onset was −273.1 ± 3.01 ms before peak gape (Fig. 6B; Table 1) and strike duration, measured as the time from strike onset to jaw closing, was 465.56 ± 26.07 ms, although ceratohyal and clavicle motion continued their motion after the jaws closed. The lower jaw depressed an average of −11.6 ± 0.85 deg with a maximum depression of −39.5 deg (Fig. 6B). Rotations of the lower jaw about the x- and y-axes and translations along all axes were within the error of the precision test.

The right ceratohyal rotated about the z-axis (depression) an average of −20.7 ± 2.01 deg, reaching peak depression 94.2 ± 30.19 ms after peak gape (Fig. 6C; Table 1). The maximum ceratohyal depression across all trials was −39.6 deg. The right ceratohyal rotated about its long-axis an average of −13.3 ± 1.50 deg, reaching peak long-axis rotation 151.4 ± 0.35 ms after peak gape (Fig. 6D; Table 1). Maximum long axis rotation of the ceratohyal was −32.5 deg.

The prey item was typically close to the jaws at strike onset, 2.0 ± 1.10 cm away from the upper jaw when the suction feeding strike began. The prey traveled an average maximum distance of 4.9 ± 0.45 cm over the suction feeding strike, from the initial prey location to the back of the oral cavity (Fig. 6F). Prey reached its maximum distance at 307.4 ± 19.4 ms after peak gape (Table 1). Maximum prey velocity was 63.3 ± 3.76 cm s^-1^ with maximum average acceleration of 2432.7 ± 366.03 cm s^-2^ (Fig. 6G/H).

The three tongue beads generally moved with the ceratohyal markers, although the distance between the markers fluctuated slightly during the strike, with the maximum distance reached at peak ceratohyal depression. The distance between right and left tongue markers ranged from 1.5 ± 0.17 cm to 1.6 ± 0.11 cm apart. The distance between right and rostral beads across both individuals ranged from 1.1 ± 0.05 cm to 1.2 ± 0.03 cm, and the left -rostral beads ranged from 1.5 ± 0.56 cm to 1.7 ± 0.59 cm apart throughout the suction feeding strike (Table S2).

### Skeletal kinematics of the clavicle and cranial rib, and muscle shortening

The clavicles and cranial rib retracted substantially in the three strikes in which they were marked (Fig. 8, Table S3). In these trials, peak ceratohyal depression (−42.3 ± 5.3 deg) was followed by peak clavicle retraction (−22.4 ± 3.3 deg), and subsequently by peak cranial rib retraction (−7.7 ± 1.8 deg), consistent with an anterior-to-posterior wave of expansion. The ceratohyal rotated internally about its long-axis −39.1 ± 3.8 deg, nearly as much as it depressed (Fig. 8; Table S3), as seen in the other trials (Fig. 6). In contrast, the clavicle performed little, long-axis rotation (4.3 ± 1.5 deg; external rotation; Fig. 8) and flaring (4.6 ± 0.5 deg; adduction).

The rectus cervicis muscle and the costoclavicular portion of the hypaxial muscles (CCH) both shortened during buccal expansion. The rectus cervicis started shortening earlier than the CCH, reaching peak strain (−10.7 ± 1.0 % Li) slightly before peak ceratohyal depression (Fig. 8, Table S3). Then, as the ceratohyals elevated, the rectus cervicis lengthened past its initial length, reaching a mean minimum strain of 14.0 ± 2.6 % Li. The CCH reached peak strain (−16.4 ± 2.6 % Li) slightly before peak clavicle retraction, and then lengthened to its initial length as the clavicle and cranial rib protracted.

### Volume measurements

The average time to peak volume was 61.7 ± 27.86 ms after peak gape, over 300 ms after strike onset (Fig. 6E; Table 1). The volume of the buccal endocast measured here increased to 3.8 ± 0.074 cm^3^ relative to the start of the strike (Fig. 6E). Using RCVC, we show that the RCVC of the lower jaw, ceratohyal, and clavicle varied across the suction feeding strike (Fig. 7). The lower jaw initially reached its peak positive RCVC (0.46 ± 0.1) at −246.0 ± 0.12 ms before peak gape. (Fig. 7B). Then as the ceratohyal began to depress, the lower jaw slowed as it approached peak gape (Fig. 7). The ceratohyals contributed to the remaining change in volume after peak lower jaw depression. The ceratohyals reached their peak RCVC (0.99 ± 0.04) at −6.0 ± 0.11 ms before peak gape (Fig. 7). The lower jaw closed quickly and showed a short negative RCVC. The ceratohyals showed negative volume change through the slow elevation of this skeletal element during buccal compression at the end of the suction feeding strike (Fig. 7B).

In the three additional trials for Lungfish B with the clavicle and cranial rib marked (Kaczmarek et al. 2022), we calculated the RCVC for the clavicles in addition to the other bones as described above. Since the cranial rib is surrounded by musculature and other soft tissue, we concluded the motions of this bone do not directly influence the volume change of the oral cavity. The RCVC of the lower jaw went from positive to negative contribution at the time of peak gape. Additionally, the ceratohyal and clavicle RCVC crossed the x-axis at the time of peak volume. The maximum RCVC of the lower jaw was 0.58 ± 0.16, and it reached its peak −42.67 ± 6.77 ms before peak gape. The RCVC of the clavicle reached its peak RCVC (0.55 ± 0.12) - 40.67 ± 10.41 ms before peak gape. The ceratohyal produced a maximum RCVC of 0.51 ± 0.003, reaching its peak 4.67 ± 4.05 ms after peak gape. Overall, the lower jaw contributes to the initial volume expansion, while the ceratohyal contributes to volume expansion once the lower jaw starts to elevate and until the buccal cavity reaches peak volume. The clavicle contributes to volume expansion consistently throughout the strike until maximum volume is reached.

### Interspecies comparisons

Data on the timing of suction feeding kinematics were gathered from 18 studies on a range of suction feeding species (Table S4). Most species in the survey were fed non-evasive prey (e.g., pieces of fish; Table S4). Species were chosen from all vertebrate lineages that perform suction feeding in water (i.e., chondrichthyans, actinopterygians, and sarcopterygians). The fastest suction feeder in the dataset was *Hippocampus erectus*, the lined seahorse, while the slowest species was *Lepidosiren paradoxa*, the South American lungfish. *P. annectens* and *L. paradoxa* were some of the slowest suction feeding species in our analysis. Time to peak gape averaged 273.1 ± 3.01 ms for *P. annectens* and 375 ± 135.0 ms for *L. paradoxa*. The other species with a similarly slow feeding strike is the swell shark, *Cephaloscyllium ventriosum*, which had an average time to peak gape of 310 ± 40 ms for one-year old individuals (Ferry-Graham, 1997). All other species had significantly faster times to peak gape (Fig. 9). The linear regression of time to peak gape against body length had an r^2^ value of 0.0021 and was not significant (p >0.05).

**Fig. 9.**
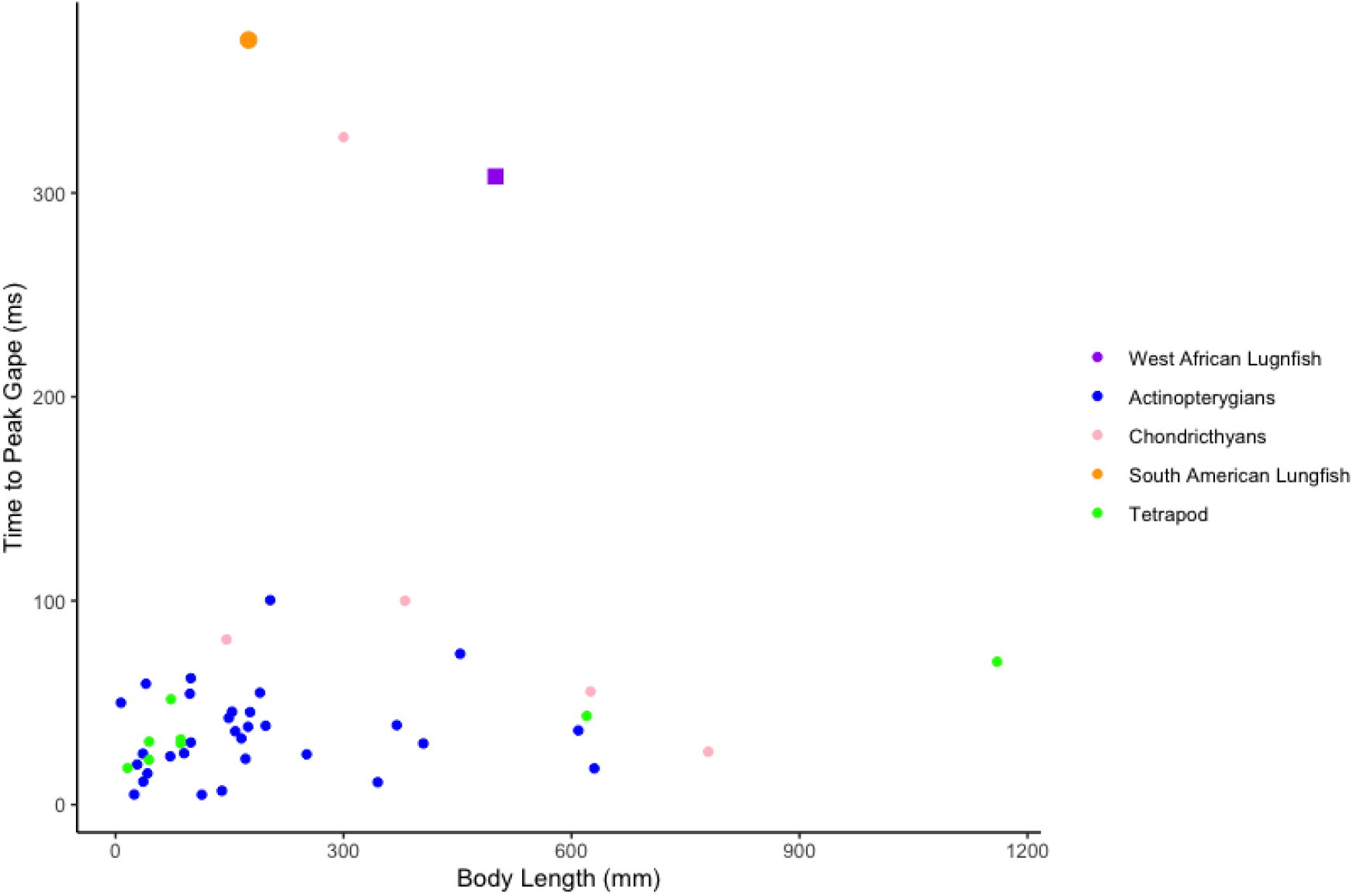
Interspecies comparison of time to peak gape. Graph of average body length (mm) vs. time to peak gape (ms) taken from a wide variety of papers recording suction feeding in water (Table S4). Purple rectangle: this study; orange circle: *Lepidosiren paradoxa* (Bemis and Lauder, 1986); pink dot: swell shark (Ferry-Graham, 1997).

## Discussion

In this study, we provide an individual-bone level analysis of the suction generation mechanism of the West African lungfish (*Protopterus annectens*), a sarcopterygian with a highly fused skull. The central conclusions of this study are that the neurocranium and lower jaw initiate the suction strike through a mechanism of flat-plate suction and that the ceratohyal undergoes complex rotations that help to drive water flow for prey transport. The volume contributions (RCVCs) of the lower jaw, ceratohyals, and clavicles show that the lower jaw initiates buccal volume change but the ceratohyals have a higher RCVC as the lower jaw approaches peak depression and the clavicles peak before maximum ceratohyal RCVC. The rectus cervicis muscle and CCH shorten during buccal expansion and lengthen throughout the strike, indicating they help drive buccal expansion. The relative timing of skull motions creates an anterior-to-posterior wave of water flow during suction feeding in *P. annectens*, similar to that of suction feeding in actinopterygians. However, we found that lungfishes capturing non-evasive prey have long duration feeding similar to that previously measured in *Lepidosiren* (Bemis and Lauder, 1986) and are among the slowest suction feeding behaviors known.

### Skeletal kinematics to create and maintain suction

The West African lungfish performs the classic aquatic suction feeding kinematic sequence seen in most bony fishes of lower jaw depression, cranial elevation, and buccal cavity expansion mediated by hyoid motion (Wainwright et al., 2015). This suggests that the basic kinematic profile of lower jaw depression followed by cranial elevation and ceratohyal depression (Fig. 6), with prey acceleration peaking at the same time as jaw depression, is a shared trait at the base of Osteichthyes (Bishop et al., 2008; Ferry et al., 2015; Lauder, 1982). During *P. annectens* suction feeding strikes, lower jaw depression occurs first and reaches its peak before maximal ceratohyal motion (Fig. 6). The relative timing of cranial elevation was variable, and in some strikes, cranial depression occurred before elevation, but peak cranial elevation was always reached after peak lower jaw depression (Fig. 6). Mean lower jaw depression was slightly less variable than cranial elevation. Although the magnitude of cranial elevation was variable, the neurocranium and lower jaw initiate the anterior-to-posterior wave of motion to move the prey into the buccal cavity.

The prey item was typically close to the oral cavity as the strike began, and then accelerated toward the mouth opening. Once the prey was transported into the mouth, it reached peak acceleration at the same time as peak lower jaw depression. The creation of sub-ambient pressure by the wide jaw pulling away from the roof of the mouth is a hydrodynamic mechanism termed flat-plate suction or leading-edge suction, used in a wide range of biological behaviors to quickly initiate fluid flow. This mechanism involves pulling two relatively flat objects apart to induce rapid flow into the vacuum created, which is characteristic of the “clap-and-fling” mechanism of rapid lift induction in flapping flight (Dickinson, 1996) and in the suction feeding mechanisms of aquatic organisms with dorsoventrally flattened heads such as gar (Lemberg et al., 2019), *Polypterus* (Whitlow et al. 2022) and giant salamanders (Heiss et al. 2013).

The prey item continues to travel with the water flow into the mouth well after the lower jaw has peaked and begun to close. The time interval during which prey acceleration and velocity are maximal indicates the period of maximal suction pressure differential between the ambient pressure and the sub-ambient buccal cavity pressure (Day et al., 2015). Peak prey velocity typically occurs at the same time as peak jaw depression and just prior to peak ceratohyal depression and clavicular retraction (Fig. 6), indicating that ceratohyal and clavicular kinematics play an important role in the movement of the prey and are key determinants of the suction feeding strike in lungfishes. The relatively low volume changes, compared to bass (Camp et al., 2015), of just 3-10 ml seen in *P. annectens* feeding on worms (Fig 6E, Fig 8C) are somewhat counter-intuitive, as the suction appears to be quite effective. We occasionally observed the prey item being expelled through the gill slit during suction feeding on smaller prey, so it is possible that there is a larger volume of water being transported, with continual flow of water through the gills. So although we don’t see large volume changes of up to 25ml similar to largemouth bass (Camp and Brainerd, 2014), lungfishes are moving sufficient water volume to enable prey capture.

Buccal expansion was generated by depression of the lower jaw, ceratohyal, clavicle, and cranial rib, along with shortening of the rectus cervicis and CCH muscles (Fig. 8A). At the caudal end of this musculoskeletal series, the cranial rib retracted and acted as an attachment site for the CCH muscle. While the cranial rib is surrounded by muscle and does not directly bound the oral cavity (Kaczmarek et al., 2022), retraction of the cranial rib pulled the CCH and clavicle caudally. The CCH shortened, likely further retracting the clavicle. Retraction of the clavicle (which defines the posterior boundary of the buccal cavity) directly contributed to oral expansion and pulled the rectus cervicis and ceratohyal caudally. Shortening of the rectus cervicis during buccal expansion likely further contributed to ceratohyal depression.

Ceratohyal motion was complex, involving both depression and long-axis rotation (Fig. 5, 6, 8), indicating that the articulation between the ceratohyals and squamosal permits motion in many degrees-of-freedom. Long-axis rotation of the ceratohyals peaked at the same time the ceratohyals reached peak depression (Fig. 6, Fig. 8B). Volume also peaked at approximately the same time as peak ceratohyal depression (Fig. 6, Fig. 8B). While the ceratohyals performed large magnitudes of internal long-axis rotation, the clavicles adducted and externally rotated by just a few degrees (Fig. 8B), likely indicating that they did not directly cause lateral expansion of the buccal cavity.

During buccal compression after the strike, the rectus cervicis and CCH had different patterns of strain. Both muscles lengthened, indicating that the associated bones upon which the muscles originate and insert spread apart as they returned to their initial positions. However, the rectus cervicis lengthened past its initial resting length, consistent with the observation that clavicle protraction lagged ceratohyal elevation, stretching the rectus cervicis as a result. Prior work on muscle activity in the rectus cervicis of *Lepidosiren* (Bemis and Lauder 1986), showed a biphasic activity pattern, with one burst early and one late in the strike, suggesting the possibility of a dual role for the muscle. If the late burst of EMG activity is present in *Protopterus* as well, that would suggest a mechanism of active lengthening and negative work, perhaps associated with the slow squeezing of water out of the buccal cavity.

We conclude that the complex motion of the ceratohyals in three dimensions, including depression, flaring and long axis rotation, enable them to achieve an elevated contribution to volume change (RCVC) than would be expected from ceratohyals with a fused symphysis. Anterior hyoid depression push the tongue ventrally, and lateral flaring increase the buccal volume rapidly. The long-axis rotation of the curved ceratohyals also serves to depress and spread their anterior tips, functioning to stretch the tongue. This pattern of complex hyoid motion, deep in the pharynx of the fish, is similar to the hyoid motions seen in *Polypterus* (Whitlow et al. 2022) and is one of the key insights gained from using XROMM to explore suction feeding. Similarly, the important roles of the highly mobile clavicles in volume expansion are insights gained only with XROMM visualization. The ceratohyals and clavicles contribute to the majority of oral cavity expansion (Fig. 7). As the oral cavity volume decreases and the hyoid begins to return to its resting position, the ceratohyals and clavicles become the main elements involved in subsequent oral cavity compression (Fig. 7), reversing their motions, pushing water out of the gill opening, and trapping the prey near the rear of the pharynx. This is demonstrated by the negative RCVC of the ceratohyals and clavicles at the time of decreasing oral cavity volume.

The extensive soft tissues of the buccal cavity play an important role in lungfish feeding. The lungfish tongue has been described as a fleshy tongue pad with little musculature for prey manipulation (Bemis, 1986), suggesting that it functions as a connection between the ceratohyals, without the ability to manipulate prey in the manner of some fishes (Sanford and Lauder, 1989), salamanders (Stinson and Deban, 2017) or mammals (Feilich et al., 2021). Our XROMM results largely support this hypothesis, with the tongue tracking the motion of the ceratohyals and helping to lower the floor of the mouth. However, our data show that the lungfish tongue increases its width in all directions as it moves with the ceratohyals, suggesting that the tongue is stretching due to the long-axis rotations of the ceratohyals. This stretching is likely facilitated by the absence of fusion of the two ceratohyals in the midline, which raises intriguing questions about the role of the material properties of soft tissues such as connective tissues, skin, and tongue in both the expansion phase and compressive phase of feeding in *Protopterus annectens*.

### How slow can you go: comparative suction feeding strategies across fishes

Lungfishes exhibit one of the slowest suction feeding strikes measured in aquatic vertebrates to capture non-evasive prey, with a relatively low volume change in the buccal cavity. Reduction of cranial elements and fusion of bones results in reduced cranial mobility in *P. annectens* (Fig. 1B, E). As such, the main sources of kinesis are the hyoid apparatus and pectoral girdle which play pivotal roles in the expansion of the oral cavity. The contribution of hyoid motion to dorso-ventral expansion is seen in other species (Camp and Brainerd, 2014), but in *P. annectens* is conducted on a relatively slow timescale. Unlike lungfishes, many other suction feeding species have extensive anterior as well as medio-lateral expansion. Lungfishes exhibit some similarities in timing patterns and kinematics, like the anterior-to-posterior wave of motion, prey processing, and flat-plate suction, but deviate in other characteristics, like gross kinematic timings.

The suction of mechanism of *P. annectens* involving an anterior-to-posterior wave of motion is similar to that of largemouth bass, *Micropterus salmoides* (Fig. 1A, D), which use suction as the main form of prey capture (Camp and Brainerd, 2014). Suction feeding of largemouth bass is initiated by lower jaw depression and elevation of the neurocranium (Camp and Brainerd, 2014). Hyoid depression and cleithrum retraction follow, further expanding the oral cavity and sucking the prey item further into the oral cavity (Camp et al., 2015; Camp and Brainerd, 2014). These fishes show the anterior-to-posterior wave of motion similar to *P.annectens*, but with lungfishes having decreased cranial elevation (Fig. 6).

*P. annectens* differs from other fish species in gross timing of the suction strike. Catfishes (Olsen et al., 2019), wrasses (Westneat, 1994, 1990) and basses (Camp and Brainerd, 2014; Wainwright et al., 2007) exhibit a typical strike duration of 50-100 ms (Table S4). This is in marked contrast with the fastest suction feeders such as seahorses that use a snap suction mechanism to feed in 1-5 ms (Van Wassenbergh et al., 2013) and frogfishes that can suck in prey larger than themselves in 6 ms (Pietsch and Grobecker, 1990). *P.annectens* takes over 250ms to complete its suction feeding strike (Fig.6). However, it is important to note that our observations of *Protopterus* feeding on evasive prey such as live feeder fish indicate that they are capable of feeding more quickly, however benthic foraging is the most commonly used mode of feeding in these fishes (Bemis, 1986).

*P. annectens* was also seen to have an extensive processing phase, similar to the kinematics of *L. paradoxa* (Bemis and Lauder, 1986) and *Cephaloscyllium ventriosum* (Ferry-Graham, 1997). Both lungfish species and the swell shark exhibit longer time to peak gape than the majority of other fish species (Fig. 9). Overall kinematics and timing patterns during suction feeding within swell sharks typically involve lower jaw depression initiating the suction event, then near maximum gape the hyoid depresses, sucking the prey further into the oral cavity (Ferry-Graham, 1997) leading to an anterior-to-posterior wave of buccal expansion during suction feeding strikes. Future research might explore the question of whether consuming non-evasive prey and having a lengthy processing phase correlates to increases in the suction strike duration. If there is a relationship, this may suggest these species that feed on hard-shelled or tough non-evasive prey also may rely on a larger volume expansion over a long period of time for successful suction feeding rather than having fast kinematics with a large increase in volume.

The flat-plate suction mechanism of *P.annectens* is similar to that of the Chinese giant salamander, *Andrias davidianus*. which uses the lower jaw as the main driver of oral cavity expansion in a mechanism thought to represent the feeding strategy of ancestral amphibians (Heiss et al., 2013). This mode of suction feeding is similar to that found in *P. annectens* in that both species generate flat-plate suction when opening the lower jaw, but it is not predicted to have been used by the ancestors of lungfishes (Clack et al., 2016). However, a difference between the feeding behaviors of these two species is that in lungfishes the hyoid has a greater influence on the generation of suction. Additionally, the flow of water during a suction feeding strike in *A. davidianus* is markedly different from that in lungfishes. The gill opening is closed so unidirectional flow is not possible in this species. Unidirectional flow of water is key to bringing prey further into the oral cavity and into the esophagus (Gibb and Ferry-Graham, 2005). Instead, *A. davidianus* has bi-directional flow (Heiss et al., 2013), creating further mechanical differences between lungfishes and salamanders. Comparisons between *P. annectens* and *A. davidianus* can help to understand some of the feeding behaviors used by tetrapodomorphs due to their phylogenetic positions on opposite sides of the water-to-land transition.

### Evolution of tetrapod suction feeding

Lungfishes are the closest living relative to tetrapods, making them important for understanding potential feeding mechanics of species in the water-to-land transition. Lungfishes are proficient at suction feeding and rely on this behavior to capture prey (Bemis, 1986; Bemis and Lauder, 1986; Otero, 2011), suggesting that having a reduced number of cranial bones and less mobile skull does not always lead to less reliance on or an inability to suction feed. Due to the reduction in skeletal elements and mobility, lungfishes may be assumed to be biting species, and fossil lungfish morphology suggests a trend towards relying more on biting as a prey acquisition strategy. However, it is assumed that fossil lungfishes captured prey through suction and primarily used their robust tooth plates to break down their prey (Clack et al., 2016). These assumption are based off living lungfishes primarily use suction feeding for prey capture (Bemis, 1986).

*Gerrothorax*, a fossil temnospondyl, is hypothesized to have used akinetic suction feeding, similar to lungfishes. *Gerrothorax* did not have cranial kinesis in its skull roof and palate, as all the sutures are rigid (Witzmann and Schoch, 2013). However, this early amphibian had hyobranchial elements (e.g., ceratohyals) that are thought to have contributed to suction feeding. The similarities between extant lungfish and a fully aquatic fossil amphibian, suggest that these fossil salamanders may have employed suction feeding similar to that shown here in *Protopterus*, mainly driven by the kinesis and depression of the hyobranchial elements.

The *Protopterus* feeding strategy shares some features, but not others, with that hypothesized for the transitionary tetrapodomorph species, *Tiktaalik roseae*. Recent research on *Tiktaalik* concluded that this transitional fish used a combination of lateral snapping and suction feeding (Lemberg et al., 2021) to capture prey. This strategy is thought to have been similar to gar, which rely on lateral biting as well as initial flat-plate suction to capture prey (Lemberg et al., 2019). The combination of suction with a lateral strike suggests the need to assess the evolutionary distribution of the switch to lateral biting in both living and fossil forms (Lemberg et al., 2021). Early amphibians with a generally flattened head also typically rely on suction feeding to capture prey in the water. We conclude that the use of a generally flattened head to generate rapid jaw depression and flat-plate suction to initiate aquatic feeding is a common feature of early sarcopterygian feeding. Further insight into lungfish feeding will help to clarify different mechanisms and strategies fishes use for suction feeding, will further our understanding of suction feeding in stem sarcopterygians, and may inform our understanding of feeding in tetrapodomorph fossils along the aquatic to terrestrial transition.

## Materials and Methods

### Specimens and sample size

Three West African lungfish (*Protopterus annectens*) (Lungfish A: total length 48.0 cm, body mass 402 g; Lungfish B: 54.0 cm, 710 g; Lungfish C: 54.0 cm, 820 g) were obtained from the aquarium industry and housed in individual aquaria (130-210 L) with both under gravel and canister filtration. All husbandry and experimental procedures followed University of Chicago IACUC Protocol 72365. Food was provided 2-3 times weekly, including earthworms, protein pellets, and occasional feeder fish. We collected 16-18 feeding strikes for each of the three individuals. After data collection, during micro-computed tomography (μCT) scanning, we discovered one of the lungfish specimens, Lungfish C, had a partial, unilateral lower jaw dislocation. Its feeding kinematics were grossly similar to those of the other two specimens, but its strikes were longer in duration and its jaw joint exhibited a reduced range of motion. Therefore, the data for the injured fish were removed from the analysis and are summarized in Table S1. The data analysis presented here is from two lungfish specimens (n= 39 feeding strikes), including 16 trials for Lungfish A, 20 trials for Lungfish B, as well as three trials from Lungfish B with markers implanted in additional bones (the left and right clavicles and the cranial rib; Kaczmarek et al., 2022).

### Surgical implantation of X-ray markers

We implanted radio-opaque tantalum markers in each lungfish specimen to facilitate quantification of cranial bone rotations using XROMM (Fig. 2D-F). Specimens were anesthetized using equal parts MS-222 and sodium bicarbonate (0.33 g/L – 0.67 g/L). About twenty-six 1.0 mm radio-opaque tantalum bead markers were placed in each lungfish in the following regions: 3 in the upper jaw (pterygoid), 3 in the lower jaw (prearticular), 3 in the right ceratohyal, 3 in the left ceratohyal, 4 in the ventral body, 1-2 in the dorsal body, 5 in the neurocranium (supraorbital), and 3 in the tongue (Fig. 2D-F). We placed three 1.0 mm markers in the tongue in a triangular shape to examine potential movements of this fleshy tongue pad (Bemis, 1986; Bemis and Lauder, 1986). In Lungfish B, additional bones were marked for three suction feeding strikes (Kaczmarek et al., 2022): three beads were implanted in each of the right and left clavicles, and hypodermic needles were used to inject one bead approximately 1.0 mm away from the left clavicle and one bead approximately 1.0 mm away from the left cranial rib. For bony elements, a hand drill was used to bore press-fit holes, while soft tissue body markers were injected using hypodermic needles. The jaws in the lungfish are extremely dense, so a High-Performance Variable Speed Rotary Tool (Dremel, Racine, WI, USA) with a 0.8 mm drill bit was used to create press-fit holes for the 1.0 mm beads in these regions. Surgeries lasted up to 90 minutes with frequent submersion in the MS-222 water.

After surgery, to visually confirm bead placements and create meshes for later animations, individuals were scanned using the University of Chicago Veterinary CT scanner (Vimago L Base version, EPICA Animal Health, Duncan, SC) lasting an additional 30 minutes. Individuals were revived in a freshwater tank and transferred to their home tank. Once fully recovered, bead implants remained intact for several months of data collection. Individuals were robust to anesthesia and handling, being able to feed and successfully capture prey within 60 minutes of being scanned.

### XROMM data collection and animation

We used the University of Chicago XROMM Facility (https://xromm.uchicago.edu/) to collect X-ray video (ProCapture VPU, Xcitex, Woburn, MA, USA) of lungfish feeding strikes on non-evasive prey (worms) implanted with 1.0 mm radio-opaque tantalum beads. Videos were filmed at 500 frames sec^-1^ in a temporary tunnel tank (585 mm L x 92 mm W x 295 mm H). X-ray videos were generated at 80 kVp or 90 kVp and 63 mA or 80 mA (right lateral view) and 95 kVp or 100 kVp and 80 mA or 100 mA (left lateral view). The three additional strikes of Lungfish B (after beads were implanted in the clavicles and cranial rib) were recorded at 150 frames sec^-1^, 75 – 80 kV and 40 mA. Individuals were kept in the smaller tank for at most an hour before being returned to their home tank. Video sequences analyzed in this study are available on the University of Chicago XROMM Data Management Portal (https://xromm.rcc.uchicago.edu/; study ID: “Lungfish_Feeding”). Sequences with a full strike (lower jaw opening to ceratohyal elevation or to the start of a processing phase) were kept and analyzed further.

XMALab (version 2.0.1) was used to undistort the X-ray images, compute the 3D camera positions, track markers, and calculate rigid body transformations (Knörlein et al., 2016). Mean marker tracking precision was calculated as the mean of the standard deviation of the unfiltered marker-to-marker distances across all bones (0.029 cm). Due to high noise in the ceratohyals, we used the polynomial fit method to refine the positions of the ceratohyal markers in XMALab. Rigid body transformations and 3D coordinates of tongue and muscle beads were filtered at 35 Hz (low-pass, butterworth filter) and exported from XMALab. The beads in the epaxial and hypaxial muscles (Fig. 2F) were used to generate the rigid body transformations for the body plane, a pseudo-rigid body (Camp and Brainerd, 2014).

A different process for filtering the tracked data and calculating rigid body transformations was used for the three strikes that were recorded from Lungfish B with clavicles and left cranial rib marked (Kaczmarek et al., 2022). For those strikes, the ‘matools’ R package (available under matools R package at https://github.com/aaronolsen) was used to smooth the 3D marker coordinates and to generate the rigid body transformations, following the XROMM workflow described in Olsen et al. (2019). The rigid body transformations for the cranial rib were produced using the marker implanted adjacent to the cranial rib as well as two virtual points placed on the cranium at the costo-cranial joint, dorsal and ventral to each other. Virtual constraints were also applied to the left ceratohyal and left clavicle (placed at the symphyses between the left and right bones). Mean marker tracking precision for these three strikes, measured as the mean of the standard deviation of the unfiltered pairwise marker-to-marker distances within all bones, was 0.12 mm, and the maximum precision error was 0.27 mm across the three trials.

Computed tomography (CT) scans (Vimago L CT Scanner, Epica, Duncan, SC, USA) and μCT scans (GE Phoenix v|tome|x 240 kv/180 kv scanner, University of Chicago Paleo-CT facility) were taken of each fish. Polygonal mesh models of each bone were created through segmentation of the scans in Horos (v.3.3.6, Horos Project, horosproject.org) or in Amira 5.5.0 (FEI Company, Hillsboro, OR, USA) through the Research Computing Center at the University of Chicago (https://rcc.uchicago.edu/). The mesh models and rigid body transformations were imported into Maya 2019 (Autodesk, San Rafael, CA, USA), creating animations of bone motion.

### Kinematic analysis

We measured the kinematic rotations and translations of the neurocranium, lower jaw, upper jaw, ceratohyal, clavicle and tongue, as well as translations of the prey during the strike. We used joint coordinate systems (JCS; Grood and Suntay, 1983) to calculate the Euler angle rotations of these bones (following the right-hand rule and zyx order of rotation) using the XROMM MayaTools package (version 2.2.3; https://bitbucket.org/xromm/xromm_mayatools/src/master/). Euler angle rotations were standardized to 0 deg by subtracting their value at the start of each strike. All kinematic variables were calculated as a mean value ± standard error.

To measure jaw depression, we placed the JCS at the posterior right jaw joint with the z-axis orthogonal to the sagittal plane and measured the rotation and translations of the lower jaw relative to the upper jaw (Fig. S1). Negative rotation about the z-axis represents depression of the lower jaw. Ceratohyal depression and long axis rotation were measured for the right ceratohyal (Fig. S1). The JCSs were placed at the posterior end of the right ceratohyal with the x-axis oriented along the long axis of the ceratohyal and with the z-axis oriented roughly parallel to the frontal plane, but approximately 30 deg offset from the sagittal plane (Fig. S1). Rotations and translations of the ceratohyal were measured relative to the neurocranium. Negative z-axis rotations measured depression of the right ceratohyal about an axis orthogonal to the ceratohyal long axis but oblique to sagittal planes. Negative rotations about the x-axis represented internal long-axis rotation of the right ceratohyal. Qualitative assessment suggested that ceratohyal movements are symmetrical, so movements of the right ceratohyal were analyzed because of higher noise values in rotation about the long-axis of the left ceratohyal (Fig. S2).

Neurocranial rotations were measured relative to the “body plane”, a pseudo-rigid body created from body marker beads (Camp and Brainerd, 2014). A JCS was paired to the body plane and the neurocranium with the z-axis oriented mediolaterally (Fig. S1). Positive z-axis rotation represented elevation. The hypaxial and epaxial markers were placed superficially, so neurocranial elevation is measuring the rotations between the body and neurocranium. To assess motion and deformation of the tongue, we measured tongue motion relative to the anterior ceratohyal and inter-marker distances among the 3 tongue markers.

Prey displacement was measured using the prey marker position in each frame relative to its initial position across the strike. The first and second derivatives of prey location were used to compute velocity and acceleration. Prey velocity and acceleration are used as a proxy for suction pressure differential, pulling the prey and a bolus of water into the mouth (Day et al., 2015). In most trials maximum prey displacement was at the posterior pharynx. However, in some trials the prey item did not reach the back of the pharynx, so maximum distance traveled does not always correspond to the back of the oral cavity. In addition, total strike duration was measured as the time from strike onset to jaw closing.

For the three strikes that were recorded from Lungfish B with the clavicles and left cranial rib marked, JCSs were placed as follows (Fig. S3; Kaczmarek et al., 2022). The JCS for the lower jaw was placed with the z-axis oriented mediolaterally (consistent with its placement for the other trials, as described above). For the ceratohyals, the z-axis of the proximal ACS was aligned to the mediolateral axis of the cranium, and the x-axis of the distal ACS was aligned along the long-axis of the ceratohyal. For the cranial rib, the z-axis of the proximal ACS was aligned to the primary axis of motion (“bucket-handle” motion, Brainerd et al., 2016) based on the joint morphology, and the x-axis of the distal ACS was aligned to the long-axis of the bone. Euler angle rotations were standardized to 0 deg by subtracting their value at the first frame of each strike. Negative z-axis rotation corresponds to depression or retraction for all bones. Negative x-axis rotation of the ceratohyals indicates internal long-axis rotation. (In Kaczmarek et al. (2022), internal long-axis rotation of the ceratohyal is a positive rotation; these values were multiplied by −1 to be consistent with this study).

### JCS Precision Study

Following the approach of Menegaz et al. (2015), we quantified the precision of the JCS motions by moving Lungfish B in front of the X-ray cameras while frozen through air, using the same X-ray settings and frame rate as the feeding trials. About 160 frames were tracked and analyzed using the XROMM workflow. The standard deviation was calculated for each degree-of-freedom of each JCS. Because a frozen specimen should have no motion at the joints, the standard deviation represents the JCS precision threshold of the XROMM method. The precision thresholds of rotations (0.80 - 1.75 degrees) and translations (−1.21 to −1.48 cm) were small relative to the motions reported for our lungfish trials.

### Muscle length changes

The lengths of the rectus cervicis and the costoclavicular portion of the hypaxial muscles (CCH) were measured in three trials from Lungfish B. The rectus cervicis originates on the clavicle and inserts on the ceratohyal, and the CCH originates on the cranial rib and inserts on the clavicle. Virtual landmarks were placed on the origin and insertion of each muscle in Maya and the distances between these landmarks were measured for each XROMM animation. Muscle strain was calculated relative to the initial muscle length (Li) measured at the first frame of lower jaw depression for each strike, with negative strain values representing muscle shortening.

### Volumetric analysis

We measured the instantaneous volume of an anterior region of the oral cavity by creating a dynamic digital endocast following the approach of Camp et al. (2020) (Fig. 3). In Maya, locators were attached to the oral surfaces within the right anterior half of the oral cavity, and their 3D coordinates over time were exported. The locator positions were then imported into MATLAB (v2020a, MathWorks, Natick, MA, USA), where, for every frame, an alpha shape was calculated from the constellation of locators, and its volume computed as double the unilateral endocast volume. The volumes were created using ‘alphashape’ function in MATLAB and an alpha value of 2. Polygonal meshes of the alpha shapes were imported into Maya to check fit (Fig. 3).

We quantified the relative contribution to oral cavity volume change (RCVC; Whitlow et al. 2022) of the ceratohyal, clavicles and lower jaw. In brief, we digitally froze individual bones relative to the neurocranium for a short time increment, on a rolling basis over the duration of a trial. Then, to calculate RCVC, we took the difference between the change in full volume and the change in frozen volume divided by the sum of the absolute values of the volume difference for each bone that was frozen. The RCVC of the *i*th bone (*RCVC_bone_i__*) to oral cavity volume change can be represented by the following equation: where for a given time increment:

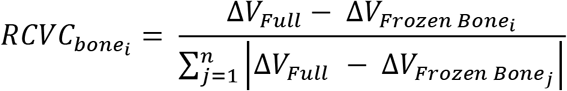

where for a given time increment:

Δ*V_Full_* is the change in endocast volume without any bones frozen and,
Δ*V_FrozenBone_i/j__* is the change in endocast volume with the *i*th or *j*th bone frozen.
*n* is the number of bones frozen (i.e., does not include the neurocranium).

We tested freeze increments of 5, 10, 15, 20, 25, and 30 frames and saw an increase in noise with lower frame counts and an over-smoothing of the data in the 25+ frame range. We concluded that a 20-frame increment (40 ms) showed the most accurate data with minimal noise for the trials recorded at 500 frames sec^-1^. For the three trials from Lungfish B that were recorded at 150 frames sec^-1^, a 12 frame increment was used.

### Interspecies comparisons

To explore comparative durations of suction feeding across species, a literature search was conducted to gather measurements of time to peak gape (ms) from actinopterygians, chondrichthyans, and salamander species, as well as the prey type (evasive vs non-evasive) in each study. If body length was not provided, we estimated the variable based off scaled figures (Table S4). We ran a linear regression using body length (mm) vs. time to peak gape (ms) in RStudio 2022 (RStudio: Integrated Development for R. RStudio, PBC, Boston, MA, USA).

## Competing Interests

The authors declare no competing interests.

## Acknowledgments

We thank Ariel Camp for sharing the endocast volume protocol, and Nick Gidmark and Nicolai Konow for implanting the first marker sets in the lungfishes, and Andrew George and Chloe Nash for help with lungfish maintenance. We are also grateful to the staff at the Animal Resource Center at the University of Chicago who helped with animal care and use of the veterinary CT scanner. Finally, we thank Neil Shubin and members of his laboratory for lending the chemicals necessary to PMA stain our specimens, and we acknowledge the efforts of our reviewers. This is UChicago XROMM Facility publication #12.

## Funding

This work was supported by the National Science Foundation (DBI 1338066 to C.F.R, DEB 1541547 to M.W.W., DGE 1644760 to E.B.K., IOS 1655756 to Elizabeth L.Brainerd, and OAC 1626552 to Hakizaumwami B. Runehsha], the Bushnell Research and Education Fund [E.B.K], and the University of Chicago [S.M.G.]. Funding for the UChicago XROMM Facility was provided by National Science Foundation Major Research Instrumentation Grants MRI 1338036 and 1626552.

## Notes

https://figshare.com/articles/journal_contribution/Supplemental_Lungfish_Feeding_Files/19912597

### Competing Interest Statement

The authors have declared no competing interest.

### Summary of Updates

We have made a number of changes to the manuscript in response to the review, including shortening and clarifying the discussion, presenting data on the tongue deformation differently, and a number of smaller changes.

