## Supplementary Figures for "Suction feeding of West African lungfish (*Protopterus annectens*): An XROMM analysis of jaw mechanics, cranial kinesis, and hyoid mobility"

**Supplementary Videos:** [**https://figshare.com/articles/journal_contribution/Supplemental_Lungfish_Feeding_Files/19912597**](https://figshare.com/articles/journal_contribution/Supplemental_Lungfish_Feeding_Files/19912597)

**Video S1:** Representative X-ray video of a suction feeding strike in a West African Lungfish (*Protopterus annectens*).

**Video S2:** Representative suction feeding animation of a West African Lungfish (*Protopterus annectens*) with the X-ray videos in the background. Blue—upper jaw; green—lower jaw; yellow—neurocranium; red—left ceratohyal; purple—right ceratohyal; orange circle—prey item.

**Supplementary Figures:**

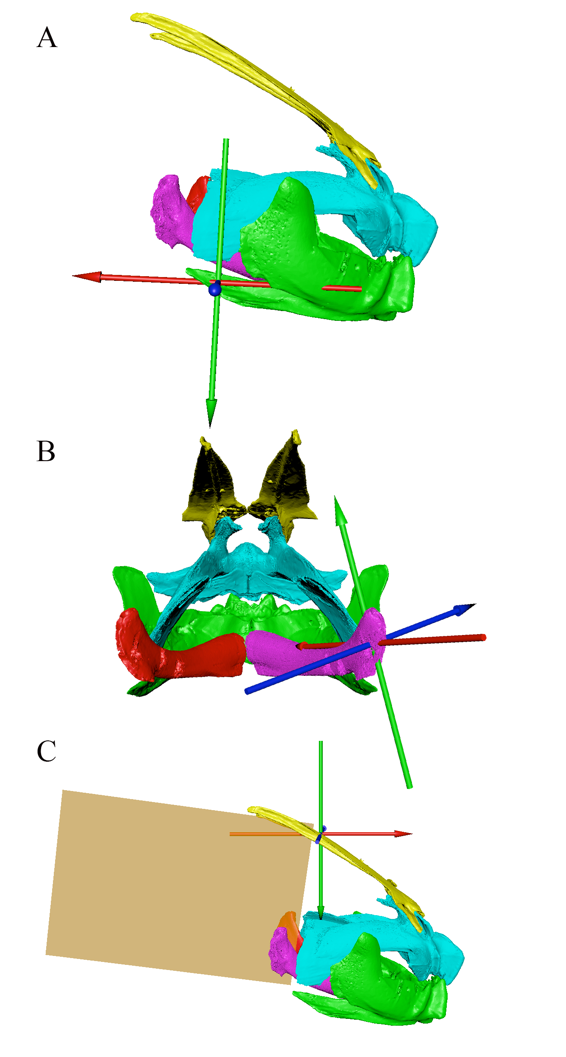

**Figure S1: Joint coordinate axis system placement in the lungfish skull and body plane.** Joint coordinate axis (JCS) placements for a representative individual for the (A) jaw joint, (B) ceratohyal joint, and (C) centrum of the neurocranium.

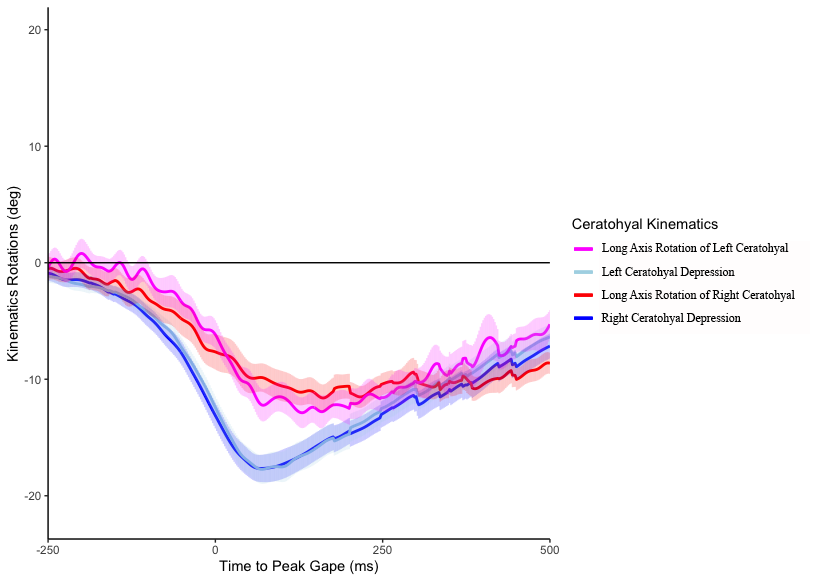

**Figure S2:** **Kinematic plot of both ceratohyals averaged across both lungfish.** The left and right ceratohyals mirror each other in their movements, but there is higher noise in the long axis rotation of the left ceratohyal due to the linearity of the implanted beads.

**Figure S3: Proximal and distal anatomical coordinate systems used for the three strikes from lungfish B in which the clavicles and cranial rib were marked.** Dorsal (A), ventral (B), lateral (C), and caudal (D) views of the placement of the proximal ACSs for the lower jaw, left and right ceratohyals, left and right clavicles, and left cranial rib. Dorsal (E), ventral (F), lateral (G), and caudal (H) views of the placement of the distal ACSs for the lower jaw, left and right ceratohyals, left and right clavicles, and left cranial rib. The motion of these ACSs was driven by their respective bones (i.e. they were parented to the bone of interest). The rotations about these JCSs are shown in Fig. 9. Note that in those figures, rotations were zeroed to the start of the behavior. Modified from Kaczmarek et al. (2022).

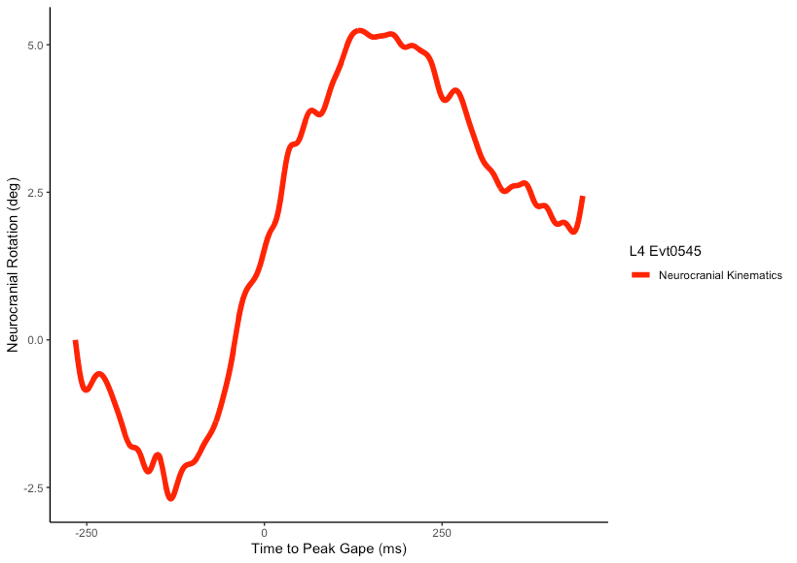

**Figure S4: Representative cranial kinematic plot.** Cranial rotations are plotted to show an example of cranial depression preceding cranial elevation.

**Supplementary Tables:**

**Table S1:** Kinematic variables of the timings and rotations during a suction feeding strike of Lungfish C, the fish with a damaged jaw joint.

| **Variables** | **Lungfish C** |
| --- | --- |
| Time to Peak Gape (ms) | 321.3 ± 183.7 |
| Jaw Depression (degrees) | 14.3 ± 3.0 |
| Ceratohyal Depression (degrees) | 28.1 ± 6.7 |
| Ceratohyal Long Axis Rotation (degrees) | 24.5 ±5.1 |
| Volume (cm^3^) | 10.0 ± 2.2 |
| Cranial Elevation (degrees) | 4.1 ± 4.8 |

**Table S2:** Tongue marker distances of maximum distance and minimum distance reached for each individual lungfish.

| **Variables (unit)** | **Lungfish A** | **Lungfish B** | **Combined** |
| --- | --- | --- | --- |
| Maximum Distance Between Middle-Right Markers (cm) | 1.2 ± 0.10 | 1.2 ± 0.31 | 1.2 ± 0.03 |
| Minimum Distance Between Middle-Right Markers (cm) | 1.1 ± 0.11 | 1.1 ± 0.32 | 1.1 ± 0.05 |
| Maximum Distance Between Middle-Left Markers (cm) | 1.1 ± 0.03 | 2.2 ± 0.33 | 1.7 ± 0.59 |
| Minimum Distance Between Middle-Left Markers (cm) | 0.9 ± 0.07 | 2.1 ± 0.30 | 1.5 ± 0.56 |
| Maximum Distance Between Right-Left Markers (cm) | 1.5 ± 0.28 | 1.7 ± 0.20 | 1.6 ± 0.11 |
| Minimum Distance Between Right-Left Markers (cm) | 1.4 ± 0.19 | 1.6 ± 0.18 | 1.5 ± 0.17 |

**Table S3.** Average kinematic variables across strikes from lungfish B in which the clavicles and cranial rib were marked. Timings are relative to the start of the suction feeding strike. Values are given as mean $\pm$ SE.

| **Variables** | **Lungfish B**  **(n = 3)** |
| --- | --- |
| Peak Jaw Depression (deg) | -14.0 ± 3.3 |
| Time to Peak Gape (ms) | 175.6 ± 50.2 |
| Peak Ceratohyal Depression (deg) | -42.6 ± 5.2 |
| Time to Peak Ceratohyal Depression (ms) | 293.3 ± 57.7 |
| Peak Ceratohyal Long-Axis Rotation (deg) | -39.1 ± 3.8 |
| Time to Peak Ceratohyal Long-Axis Rotation (ms) | 362.2 ± 89.2 |
| Peak Clavicle Retraction (deg) | -23.5 ± 4.1 |
| Time to Peak Clavicle Retraction (ms) | 344.4 ± 75.5 |
| Peak Cranial Rib Retraction (deg) | -7.7 ± 1.8 |
| Time to Peak Cranial Rib Retraction (ms) | 382.2 ± 97.9 |
| Peak Rectus Cervicis Strain (% L_i_) | 10.7 ± 1.0 |
| Time to Peak Rectus Cervicis Strain (% L_i_) | 240 ± 39.1 |
| Peak CCH Strain (% L_i_) | 16.4 ± 2.6 |
| Time to Peak CCH Strain (% L_i_) | 317.8 ± 53.9 |
| Peak Volume Change (mm^3^) | 8581.8 ± 1128.9 |
| Time to Peak Volume Change (ms) | 306.7 ± 56.7 |

| **Table S4:** Kinematic variables including strike timing and motion from the literature search for feeding strikes from a wide range of aquatic species. | | | | | | | | | |
| --- | --- | --- | --- | --- | --- | --- | --- | --- | --- |
| **Species** | **Type of Animal** | **Time to Peak Gape (ms)** | **Lower Jaw Depression (degrees)** | **Cranial Elevation (degrees)** | **Time to Peak Hyoid Depression (ms)** | **Total Body Length (mm)** | **Time to Peak Gape/ Body Length** | **Prey Type** | **Author** |
| Lepidosiren paradoxa | Fish | 375 ± 135.0 |  |  | 480.0± 169.0 | 175 | 2.1 | Non-elusive | Bemis and Lauder 1986 |
| Cephaloscyllium ventriosum | Shark | 327.3 ± 88.0 | 112.3 ± 19.1 | -17.3 ± 4.0 | 340.5± 78.5 | 300 | 1.1 | Non-elusive | Ferry-Graham 1997 |
| Protopterus  annectens | Fish | 308.1 ± 75.4 |  |  |  | 500 | 0.6 | Non-elusive | Gartner et al. |
| Choerodon anchoragoc | Fish | 100.3 ± 5.8 | 26.2 ±  0.2 | 4.1 ± 0.1 | 91.5 ± 33.0 | 203.5 | 0.5 | Non-elusive | Gibb and Ferry-Graham 2005 |
| Triakis semifasciata | Shark | 100.0 |  |  |  | 381 | 0.3 | Non-elusive | Motta et al. 2002 |
| Negaprion brevirostris | Shark | 81.0 |  |  |  | 146 | 0.6 | Non-elusive | Motta et al. 2002 |
| Aspius aspius | Fish | 74.0 ±  6.0 |  | 9.5 ± 1.8 | 86.0 ± 6.0 | 453.5 | 0.2 | Elusive | Wassenbergh and De Rechter 2011 |
| Andrias davidianus | Tetrapod | 70.1 ±  7.3 |  |  | 85.0 ± 21.9 | 1160 | 0.1 | Non-elusive | Heiss et al. 2013 |
| Periophthalmus barbarus | Fish | 62.0 ±  6.0 | 104.0 ± 7.8 |  |  | 99 | 0.6 | Non-elusive | Michel et al. 2014 |
| Poecilia sphenops | Fish | 59.3 ±  1.4 | 20.4 ±  0.9 | 17.2 ± 1.2 | 82.5 ± 2.7 | 40 | 1.5 | Non-elusive | Gibb and Ferry-Graham 2005 |
| Heterodontus francisci | Shark | 55.5 |  |  |  | 625 | 0.1 | Non-elusive | Motta et al. 2002 |
| Coris gaimardc | Fish | 54.89 ± 3.9 | 24.1 ±  3.6 | 4.1 ± 0.7 | 55.7 ±  10.3 | 190 | 0.3 | Non-elusive | Gibb and Ferry-Graham 2005 |
| Awaous guamensis | Fish | 54.4 ±  5.1 | 24.8 ±  2.8 | 3.1 ± 0.4 | 78.2 ±  5.7 | 97.8 | 0.6 | Non-elusive | Maie et al. 2009 |
| Tylototriton verrucosus | Tetrapod | 51.7 ± 7.6 | 52.4 ± 10.9 | 19.3 ± 4.4 | 57.1 ±  10.7 | 73 | 0.7 | Non-elusive | Heiss and Vylder 2016 |
| Anableps anableps (aquatic) | Fish | 50.0 ± 5.0 | 211.0 ± 6.0 |  |  | 7.1 | 7.0 | Non-elusive | Michel et al. 2015 |
| Lepomis macrochirus | Fish | 45.6 ± 1.8 | 39.9 ± 1.0 | 2.7 ± 0.2 |  | 153.3 | 0.3 | Non-elusive | Mehta and Wainwright 2007 |
| Oxycheilinus digrammusc | Fish | 45.4 ± 0.8 | 25.1 ± 2.3 | 5.5 ± 0.6 | 63.2 ±  0.9 | 177 | 0.3 | Non-elusive | Gibb and Ferry-Graham 2005 |
| Lepomis macrochirus | Fish | 42.5 ± 3.6 | 39.5 ± 1.7 | 9.0 ± 0.3 | 51.0 ±  3.4 | 149 | 0.3 | Non-elusive | Gibb and Ferry-Graham 2005 |
| Gambusia affinis | Fish | 39.0 ± 7.5 | 37.7±3.2 |  |  | 370 | 0.1 | Non-elusive | Ferry-Graham, Hernandez, Gibb, Pace 2010 |
| Hologymnosus doliatusc | Fish | 38.7 ± 0.3 | 16.2 ± 0.1 | 9.9 ± 6.3 | 68.5 ±  0.2 | 197.5 | 0.2 | Non-elusive | Gibb and Ferry-Graham 2005 |
| Micropterus salmoides | Fish | 38.2 ± 1.2 | 40.1 ± 1.0 | 9.5 ± 0.0 |  | 174.3 | 0.2 | Non-elusive | Mehta and Wainwright 2007 |
| Anguilla rostrata | Fish | 36.4± 2.3 | 8.2 ± 0.7 | 3.6 ± 0.4 |  | 609 | 0.1 | Non-elusive | Mehta and Wainwright 2007 |
| Novaculichthys taeniourusc | Fish | 36.0 ± 4.1 | 41.9 ± 1.8 | 4.9 ± 0.8 | 50.8 ±  5.7 | 157.5 | 0.2 | Non-elusive | Gibb and Ferry-Graham 2005 |
| Xystreurys liolepisf | Fish | 32.5 ± 4.5 | 33.1 ± 1.0 | 12.5 ± 0.5 | 69.8 ±  6.3 | 165.5 | 0.2 | Non-elusive | Gibb and Ferry-Graham 2005 |
| Lissotriton vulgaris (terrestrial stage) | Tetrapod | 32.0± 6.0 |  | 7.6 ± 4.2 | 25.0 ±  3.0 | 85.8 | 0.4 | Non-elusive | Heiss, Aerts, Van Wassenbergh 2015 |
| Ichthyosaura alpestris (metamorphed) | Tetrapod | 30.9 ± 6.4 |  | 18.6 ± 4.8 | 41.6 ±  10.6 | 44.2 | 0.7 | Non-elusive | Heiss and Grell 2019 |
| Amphilophus citrinellus | Fish | 30.5 ± 2.4 | 37.3 ± 1.3 | 8.0 ± 0.7 |  | 99 | 0.3 | Non-elusive | Mehta and Wainwright 2007 |
| Lissotriton vulgaris (aquatic stage) | Tetrapod | 30.0 ± 6.0 |  | 15.0 ± 5.0 | 25.0 ±  6.0 | 85.8 | 0.3 | Non-elusive | Heiss, Aerts, Van Wassenbergh 2015 |
| Fundulus rubrifrons | Fish | 30.0 ± 8.0 | 35.3 ± 3.4 |  |  | 405 | 0.1 | Non-elusive | Ferry-Graham, Hernandez, Gibb, Pace 2010 |
| Ginglymostoma cirratum | Shark | 26.0 ± 1.0 |  |  |  | 780 | 0.03 | Non-elusive | Motta et al. 2002 |
| Lentipes concolor | Fish | 25.2 ± 2.1 | 37.5 ± 1.7 | 7.1± 0.6 | 53.8 ±  4.5 | 90.2 | 0.3 | Non-elusive | Maie et al. 2009 |
| Kryptolebias marmoratus | Fish | 25.0 $\pm$ 5.0 |  |  |  | 36 | 0.7 | Non-elusive | Ferry-Graham, Gibb, Hernandez 2008 |
| Syngnathus leptorhynchus | Fish | 24.7 ±  9.5 | 14.5 ± 2.6 | 25.1 ± 1.4 | 36.7 ±  2.3 | 251.5 | 0.1 | Non-elusive | Gibb and Ferry-Graham 2005 |
| Heniochus acuminatusb | Fish | 23.7 ± 6.9 | 19.6 ± 1.0 | 4.2 ± 0.3 | 50.7 ±  2.1 | 72 | 0.3 | Non-elusive | Gibb and Ferry-Graham 2005 |
| Pleuronichthys verticalisg | Fish | 22.5 ± 2.7 | 33.1 ± 0.6 | 27.6 ± 0.9 | 103.4±10.3 | 171 | 0.1 | Non-elusive | Gibb and Ferry-Graham 2005 |
| Ichthyosaura alpestris (paedeomorph) | Tetrapod | 22.0 ± 5.6 |  | 16.6 ± 4.8 | 38.0 ±  9.8 | 44 | 0.5 | Non-elusive | Heiss and Grell 2019 |
| Clinocottus analis | Fish | 19.7 |  |  | 25.0 | 28.5 | 0.7 | Non-elusive | Cook 1996 |
| Salamandra salamandra | Tetrapod | 18.0 ± 7.4 | 35.0 ± 2.4 | 31.5 ±  2.0 | 24.6 ±  1.1 | 15.8 | 1.1 | Non-elusive | Reilly 1995 |
| Chaetodon xanthurus | Fish | 17.8 ± 3.7 | 19.8 ± 1.4 | 4.3 ±  0.7 | 18.8 ±  3.1 | 630 | 0.03 | Non-elusive | Gibb and Ferry-Graham 2005 |
| Betta splendens | Fish | 15.3 ± 2.5 | 42.1 ± 1.6 | 21.5 ± 0.7 | 37.0 ±  11.8 | 42 | 0.4 | Non-elusive | Gibb and Ferry-Graham 2005 |
| Danio rerio | Fish | 11.3 ± 1.2 | 41.7 ± 1.4 | 21.9 ±  0.8 | 15.3 ±  1.5 | 36.5 | 0.3 | Non-elusive | Gibb and Ferry-Graham 2005 |
| Belonesox belizanus | Fish | 11.0 ± 1.0 | 89.7 ± 4.1 |  |  | 345 | 0.03 | Non-elusive | Ferry-Graham, Hernandez, Gibb, Pace 2010 |
| Syngnathus floridae | Fish | 6.8 ±  2.8 |  | 29.2 ±  8.5 | 6.1 ±  2.0 | 140 | 0.05 | Non-elusive | Bergert and Wainwright 1997 |
| Syngnathus leptorhynchus | Fish | 5.0 |  |  |  | 24.55 | 0.2 | Non-elusive | Ferry-Graham, Gibb, Hernandez 2008 |
| Hippocampus erectus | Fish | 4.9 ±  1.8 |  | 29.1 ±  6.9 | 4.7 ±  1.3 | 113.5 | 0.04 | Non-elusive | Bergert and Wainwright 1997 |
